# Loop Extrusion Accelerates Long-Range Enhancer-Promoter Searches in Living Embryos

**DOI:** 10.64898/2026.02.17.706355

**Authors:** Pavan Choppakatla, Aleena L. Patel, Tohn Borjigin, Tee Udomlumleart, Jie Hu, Thomas Gregor, Alistair N. Boettiger, Michael S. Levine

**Affiliations:** Lewis-Sigler Institute for Integrative Genomics, Princeton University, NJ, USA; Department of Developmental Biology, Stanford University, Stanford, CA, USA; Department of Chemical and Biological Engineering, Princeton University, NJ, USA; Department of Genetics, Stanford University, Stanford, CA, USA; Department of Developmental and Stem Cell Biology, CNRS UMR3738 Paris Cité, Institut Pasteur, 75015 Paris, France; Joseph Henry Laboratories of Physics, Princeton University, Princeton, NJ 08544, USA; Department of Molecular Biology, Princeton University, NJ, USA

## Abstract

Long-range gene regulation underlies a variety of human developmental disorders including Cornelia de Lange syndrome^1^ and polydactyly^2^. Cohesin-mediated loop extrusion and tether-like elements are two major mechanisms implicated in fostering long-range enhancer-promoter (E-P) contacts^3^. However, our understanding of the contributions of these mechanisms to the kinetics of E-P interactions is limited. Here we employ a combination of quantitative single-cell imaging, genetic manipulations and polymer simulations to examine the interplay of cohesin-mediated loop extrusion and tethering elements^4,5^ in the timely activation of gene expression in living *Drosophila* embryos. Depletion of NIPBL or deletion of an enhancer-proximal CTCF anchor element reduced expression without changing the duration of individual transcriptional bursts. Genetic epistasis experiments recapitulated polymer simulations predicting complementation of tether deletions by augmenting the stability of cohesin via reduced levels of WAPL^6^. We propose a “scan and snag” model whereby directional cohesin-driven enhancer scanning promotes diffusion-mediated tethering interactions to produce successful E-P contacts and transcriptional activation. We discuss how modulating cohesin stability and the “stickiness” of looping factors^7–10^ can fine-tune the levels and timing of gene expression in mammalian developmental and disease processes.

## INTRODUCTION

Animal development is orchestrated by transcriptional enhancers that control the spatial and temporal dynamics of gene activity. Many work over distances of tens to hundreds of kilobases^11–13^. How information flows across such large intervals, and how long this takes, is poorly understood. Recent investigations in vertebrates have highlighted a role for the cohesin complex^14–21^.

Cohesin is believed to catalyze loop extrusion, the bidirectional reeling of chromatin to bring together distant sequences^22^. This process is dependent on the accessory protein complex, NIPBL-MAU2, and has been reconstituted *in vitro*^23^. In vertebrate genomes, loop extrusion is constrained by CTCF^24,25^ and can explain the formation of ‘topologically associated domains’ (TADs)^26–31^. TAD boundaries restrict regulatory information, and their loss can cause misexpression^2,32–35^.

The role of cohesin in regulating long-range enhancer-promoter (E-P) contacts within individual TADs is less well understood. Genes regulated by remote enhancers are preferentially disrupted by reductions or loss of NIPBL or cohesin whereas proximal enhancers mapping within ~50 kb of their target promoters are largely resistant^14,16–18^. Despite evidence for dynamic cohesin loops^15,36,37^, the mechanisms of forming E-P contacts remain uncertain. This is especially true for ultra-long-range interactions at critical developmental control genes such as *Sox9*^38^ and *Shh*^21,39^.

Cohesin-independent mechanisms have also been implicated in long-range gene regulation. In *Drosophila*, pairs of tethering elements have been shown to form high-frequency associations in Micro-C contact maps. There is evidence that these associations are mediated by homotypic interactions^40,41^. Tether-tether interactions can span large distances and bypass TAD boundaries^4,40,42,43^. Tether-like elements that confer long-distance activity to enhancers have also been identified for distal limb enhancers in vertebrates, and there is evidence that LIM-domain proteins LDB1 and LHX2 play a role in this activity^9,44,45^.

To investigate the interplay of tethers and cohesin in dynamic E-P contacts we used a combination of *cis* and *trans* perturbations in living *Drosophila* embryos to measure long-range E-P interactions over a distance of 250 kb. Using single-cell live imaging of transcriptional bursts, we find that cohesin and tethers both increase the speed and probability of successful transcription. We propose a “scan and catch” model whereby an enhancer-adjacent CTCF binding site anchors cohesin to work in concert with tethering elements for the timely activation of gene expression by remote enhancers. We discuss the implications of these findings with respect to long-range gene regulation in mammalian developmental and disease processes.

## RESULTS

### *Drosophila* NIPBL enables long-distance gene regulation

To investigate the potential roles of Nipped-B in long-range gene regulation in *Drosophila*, we generated a homozygous fly line with the mini-AID degron tag^46^ at the 3’ of the endogenous *Nipped-B* allele (**Fig. 1a**, hereafter referred to as NIPBL). Upon expression of the plant E3-ubiquitin ligase variant OsTIR(F74G)^46^ and the addition of the egg-permeable auxin analog, 5-PHH-IAA-AM^47^, we obtained rapid degradation of >90% of the endogenous NIPBL protein (**Fig. 1b, 1c, Supplementary Fig. 1a, 1b**). NIPBL-depleted embryos appear to proceed successfully through multiple mitoses (**Supplementary Fig. 1c**). However, these embryos fail to hatch even after a short early pulse of the auxin analog, strongly suggesting a critical role of NIPBL in developmental gene regulation (**Supplementary Fig. 1d**).

**Fig. 1.**
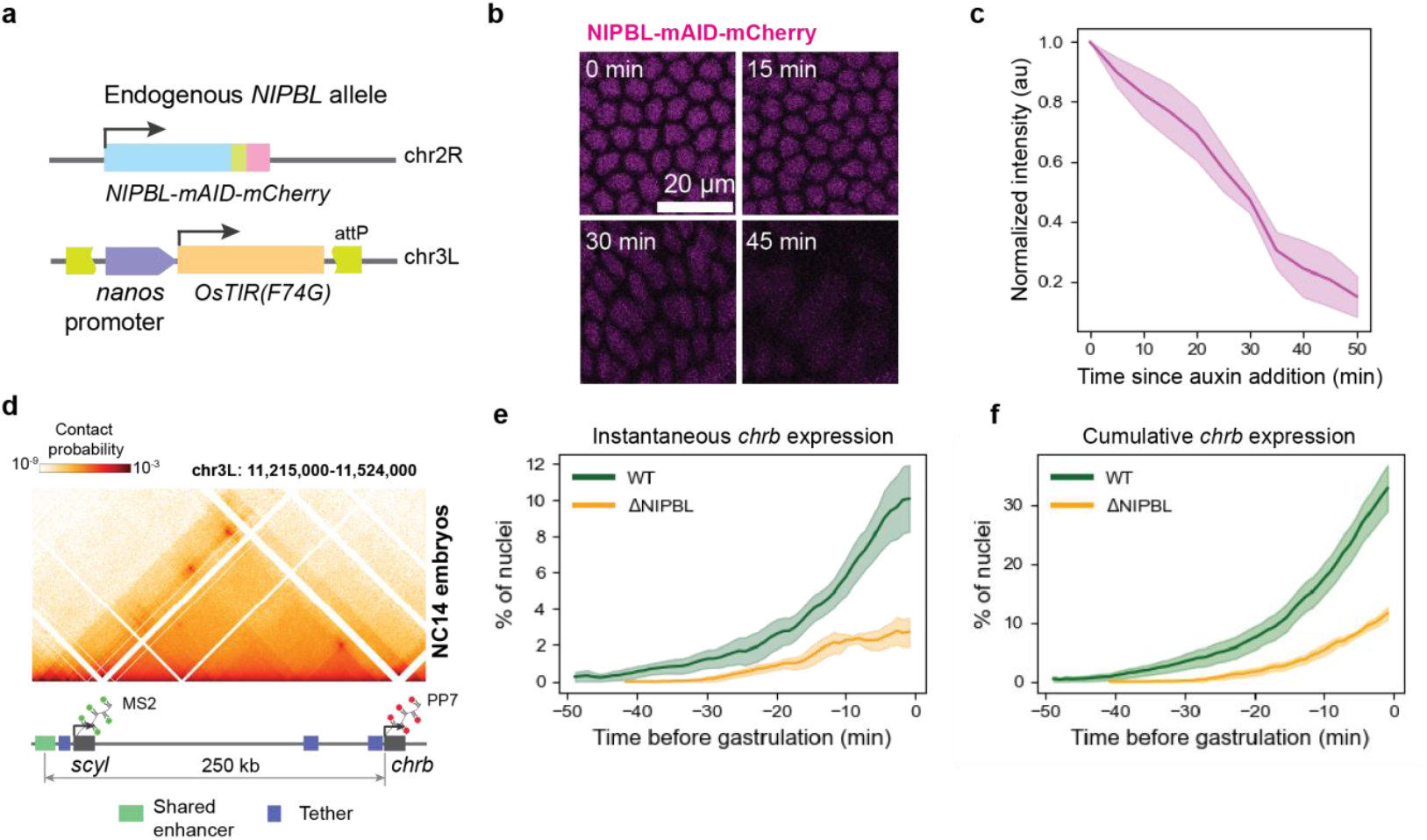
*Drosophila* NIPBL enables distant enhancer activity. **a**, Schematic of the Auxin-Inducible Degron (AID) system, showing the endogenous tagging of the NIPBL allele and the OsTIR1 transgene. **b**, Time-lapse snapshots of a representative embryo demonstrating NIPBL depletion following auxin addition at T=0. **c**, Degradation kinetics showing background-corrected and normalized NIPBL-AID-mCherry intensity after treatment with 50 μM auxin. Data represent mean +/-s.e.m. **d**, Micro-C contact map (800 bp resolution) of the *scyl-chrb* locus in NC14 embryos showing the strong contacts between tethering elements and the location of the shared dorsal band enhancer^4^. **e**, Percentage of nuclei (with active enhancer) with an active PP7 transcriptional burst in WT (WT *scyl/chrb* allele + DMSO, n=8 embryos) and ΔNIPBL (WT *scyl/chrb* allele + Auxin, n=6 embryos) embryos. Expression data were aligned at the start of gastrulation. Data plotted is mean +/-s.e.m. **f**, Cumulative percentage of nuclei showing at least one *chrb* transcriptional burst by the indicated time. Embryos and alignment are the same as in **e**.

The *charybde* (*chrb*) gene is regulated by a dorsal ectoderm enhancer that is located 250 kb away, one of the longest E-P interactions in the fly embryo^4,11^ (**Fig. 1d**). This enhancer also activates the nearby paralog *scylla* (*scyl*). The *scyl-chrb* locus thereby offers an ideal natural system to study the flow of information at long-range, as *scyl* is activated early and records the activation by the enhancer, while the subsequent activation of *chrb* provides a real-time measure of how long it takes for this regulatory information to travel a further 250 kb to the *chrb* promoter.

To examine the impact of NIPBL depletion on *chrb* transcription, we combined auxin-induced NIPBL-depletion with live-imaging of nascent transcripts using MS2 and PP7 stem loops inserted into the endogenous *scyl* and *chrb* genes, respectively^4^ (**Fig. 1d**). Expression of *scyl* was not strongly affected by NIPBL depletion due to proximity of the shared enhancer (**Supplementary Fig. 2a, 2b**). By contrast, *chrb* expression showed marked reductions in signal intensities (**Supplementary Fig. 2c, 2d**) and in the number of expressing nuclei (**Fig. 1e, 1f**). NIPBL-depleted embryos also took substantially longer than untreated embryos to reach the same fraction of activated nuclei (**Figure 1f**). There was no loss of *chrb* expression in auxin-treated control embryos lacking the OsTIR E3 ligase (**Supplementary Fig. 2e**). These observations suggest that *Drosophila* NIPBL is similar to vertebrate NIPBL in enabling long-distance enhancer-mediated regulation ^14,17,20,48^.

To determine whether cohesin is required to maintain stable contacts during bursting, we analyzed PP7 transcription traces in single nuclei (**Supplementary Fig. 2f**). Since many bursts occur at the end of the imaging interval and are artificially truncated, we modeled the bursts with an accelerated-failure time model and obtained a prediction of the true transcription profiles. NIPBL-depletion resulted in no significant changes in the modeled burst ON and OFF durations (**Supplementary Fig. 2g, 2h**) or instantaneous burst intensities (**Supplementary Fig. 2i**). The clear delay in *chrb* activation, strong reduction in the number of *chrb* expressing nuclei, and unaffected burst durations suggest that the primary role of cohesin is to accelerate long-distance E-P searches.

### CTCF-anchored loop extrusion facilitates E-P scanning

Anchoring cohesin near the enhancer or promoter is predicted to produce directional scanning of one element towards the other. In mammals, CTCF has been shown to stall cohesin extrusion^24,49,50^ and preferentially lie at the base of cohesin-dependent DNA loops^19,49–52^. We therefore deleted the enhancer-adjacent CTCF element (ΔCBS) upstream of *scyl*^4^ (**Fig. 2a**) and observed a dramatic delay and reduction in *chrb* expression (**Fig. 2b, Supplementary Fig. 3a, 3b**), without a significant change in the modeled ON or OFF durations (**Supplementary Fig. 3c, 3d**). These results strongly suggest a role for cohesin mediated loop extrusion in promoting long distance E-P contacts at the *scyl/chrb* locus. It is not currently possible to visualize and track single cohesin molecules and their extruded DNA at defined *loci* in living embryos. We therefore employed polymer simulations to determine whether cohesin might function as a DNA loop extruding motor, as suggested by studies in vertebrate systems^23,26,31^. We modeled DNA as a flexible polymer and cohesin as a factor that loads nonspecifically across the polymer and extrudes DNA bidirectionally^26^ (**Supplementary Fig. 3e**). Loop extrusion was limited by cohesin stability, which was determined by WAPL concentration^6,53^.

**Fig. 2.**
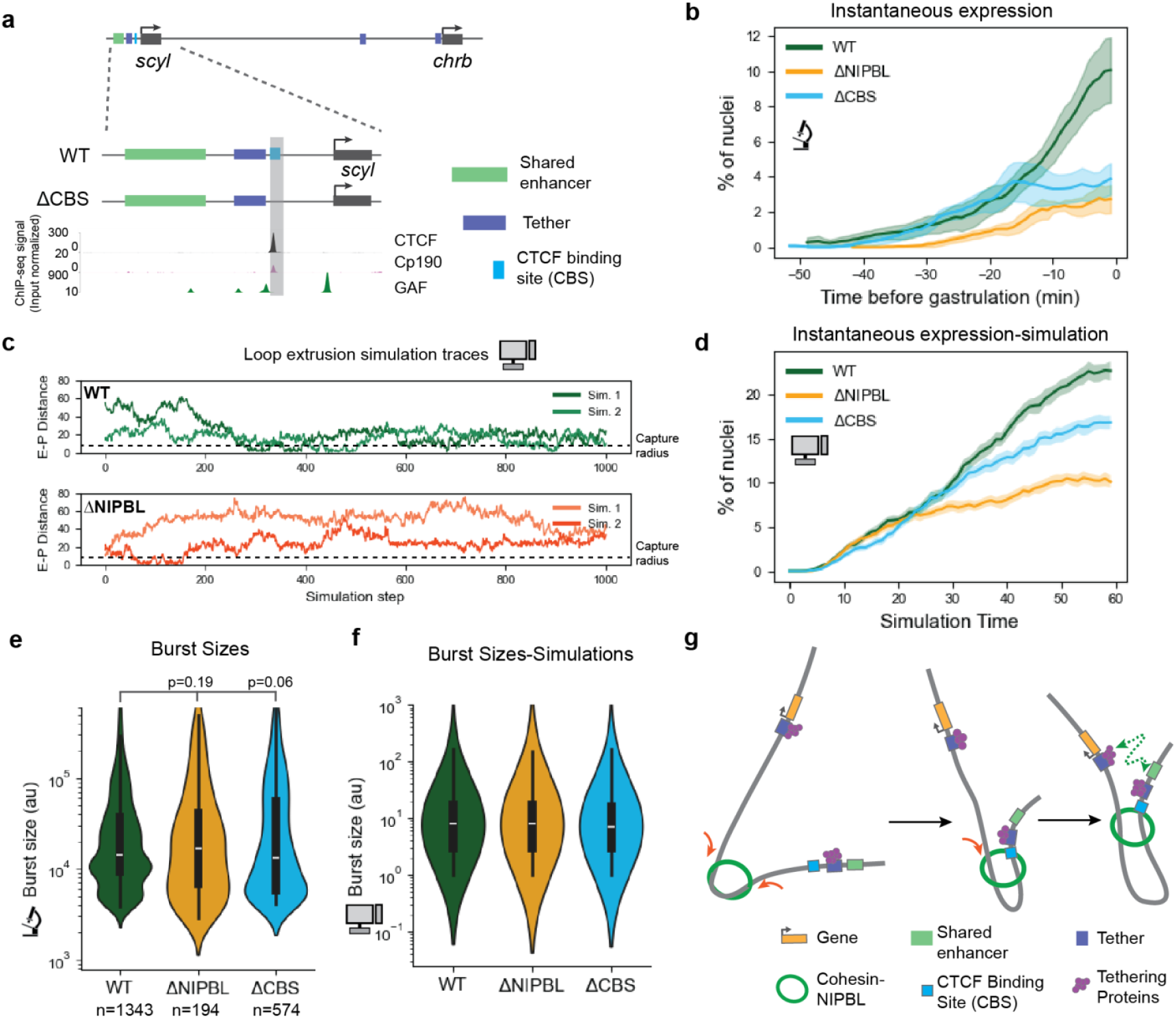
Directed loop extrusion drives *chrb* expression. **a**, Genomic organization of *cis*-regulatory elements at *scyl* (chr3L: 11,236,000–11,257,000). Tracks show ChIP-seq profiles for CTCF, Cp190 and GAGA-associated factor (GAF) in NC14 embryos^61^. The ΔCBS allele indicates the specific deletion of the CTCF-binding site. **b**, Percentage of nuclei (with active enhancer) exhibiting at least one *chrb* transcriptional burst at the indicated time. Data for WT and ΔNIPBL-depleted embryos (from **Fig. 1f**) are compared to the ΔCBS allele (n=7 embryos). Data plotted is mean +/-s.e.m. **c**, Representative polymer simulation traces comparing **WT** (active loop extrusion) with a **ΔNIPBL** (passive diffusion only). The “capture radius” defines the maximum distance from the *chrb* promoter where enhancer contact triggers expression. Distance is normalized to the size of monomer. **d**, Simulation-derived kinetics showing percentage of nuclei (% of simulations) with at least one burst at the indicated time (simulation step) for WT, ΔNIPBL (no loop extrusion) and ΔCBS (loop anchor deletion). Data show is mean and 95% C.I. **e**, Distribution of transcriptional burst sizes in the indicated conditions (embryos from **b**). Boxplots indicate the median (white line), interquartile range (IQR; black box), and 1.5 x IQR (whiskers). P-values were calculated using a two-sided Mann-Whitney U-test; n indicates the total number of bursts analyzed. **f**, Predicted transcriptional burst sizes from the polymer simulations in **d**, formatted as in **e. g**, Model of CTCF-anchored loop extrusion. Cohesin-mediated extrusion (red arrows) reduces the effective distance between the enhancer and promoter (E-P), allowing passive diffusion to facilitate tether-tether interactions.

As extrusion is dependent on NIPBL^23^, its depletion was modeled by diffusive polymer motion without extrusion. When cohesin encountered monomers occupied by CTCF, extrusion was stalled in that direction but nonetheless continued in the other direction. Finally, tether-tether interactions were captured as adhesive interactions, between pairs of “sticky” monomers. We coupled this widely used model of chromatin folding^26,54^ to a simple model of transcription whereby burst probabilities increase with diminishing E-P distances^55,56^.

Purely diffusive motion (ΔNIPBL) resulted in rare E-P contacts, whereas the addition of loop extrusion greatly increased the frequency of these contacts (**Fig. 2c**). This increase depended on the average cohesin loop size and the strength of the tethering interactions (set by the number of tether monomers) in the model (**Supplementary Fig. 3f, g**). Average cohesin loop sizes of 100 kb were sufficient to reproduce the ~70% reduction in the normal number of expressing nuclei in the absence of loop extrusion (**Fig. 2d, Supplementary Fig. 3f**). These observations are consistent with the importance of loop extrusion over passive diffusion in long-distance E-P search^15^.

Simulations of loop extrusion, in which the CTCF site near the enhancer was removed, recapitulated the delay in onset and reduced number of expressing nuclei (**Fig. 2d**). Depletion of NIPBL or deletion of the distal CTCF anchor delayed activation but did not alter burst sizes both experimentally and in the simulations since the *chrb* promoter region lacks a loop extrusion anchor (**Fig. 2e, 2f**). These observations suggest that E-P search times are accelerated by unidirectional loop extrusion (**Fig. 2g**), consistent with the view that NIPBL acts as an extrusive motor in *Drosophila*, as seen in vertebrates.

### Tethers augment E-P independently of loop extrusion communication

Our model indicates that the average cohesin loop size is smaller than the E-P distance. This means that most cohesin loops do not enable direct E-P contacts. As tethers have been shown to play a major role in *chrb* expression^4^, we measured their contributions by deleting the tether adjacent to the distal enhancer (ΔTether **Fig. 3a**). This resulted in a significant loss of *chrb* transcription, both in terms of the fraction of expressing nuclei and total expression levels (**Fig. 3b, 3c, Supplementary Fig. 4a**). There is only a modest reduction in burst sizes, as seen previously^3^ (**Fig. 3d**). There were also no significant changes in the average burst intensities (**Supplementary Fig. 4b**) or modeled ON or OFF durations (**Supplementary Fig. 4c, 4d**).

**Fig. 3.**
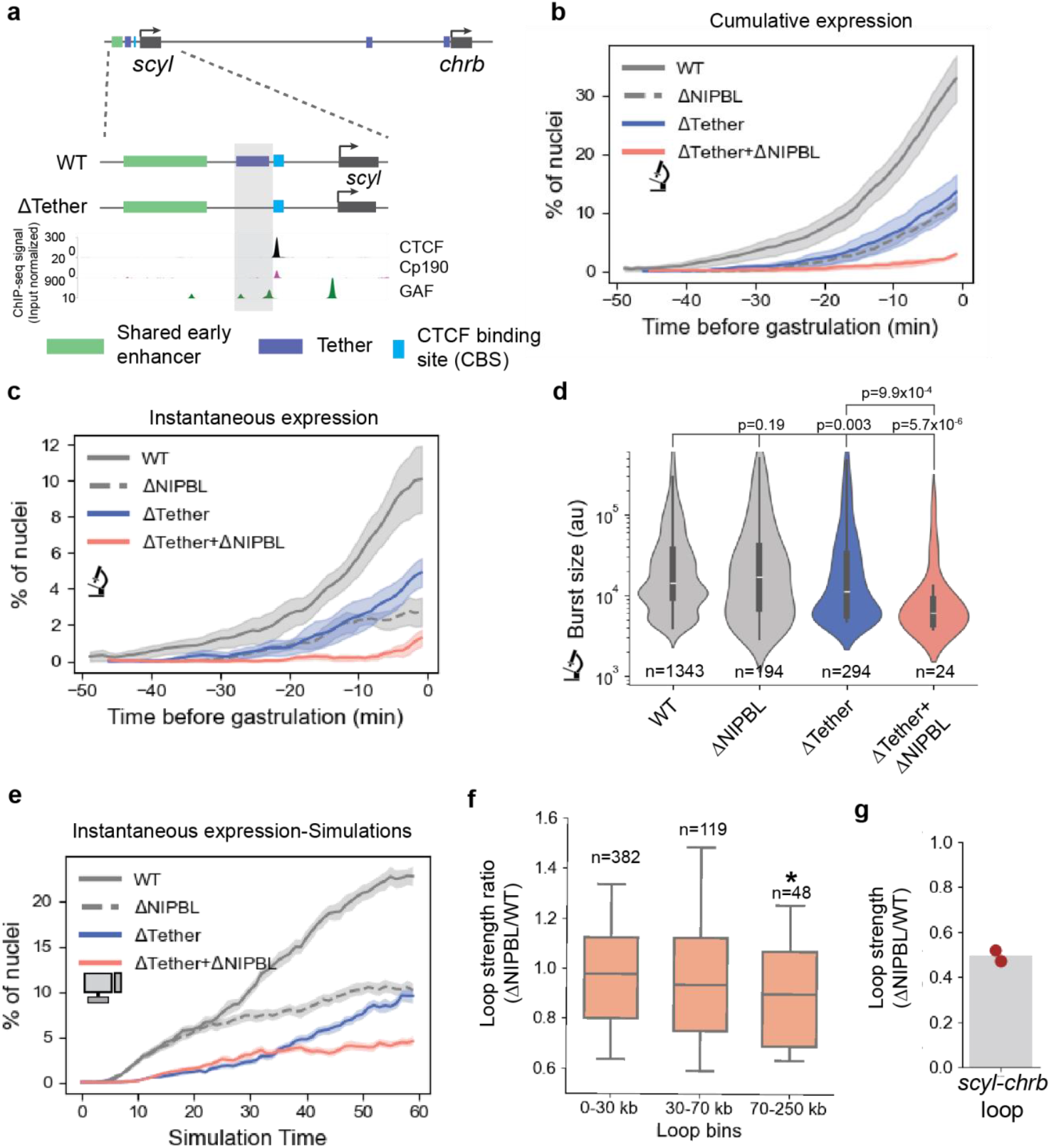
Loop extrusion and tethers cooperate to enable long-range enhancer function. **a**, Schematic showing the organization of *cis-*regulatory elements near *scyl* in WT and the distal tether deletion in ΔTether alleles. ChIP-seq tracks of CTCF, Cp190 and GAGA-associated factor (GAF) are from NC14 embryos^61^. **b**, Percentage of nuclei (with active enhancer) with an active PP7 transcriptional burst. WT and ΔNIPBL are the same as in **Fig. 1e** were compared to ΔTether (ΔTether *scyl/chrb* allele, n=6 embryos) and ΔTether +ΔNIPBL (ΔTether *scyl/chrb* allele + Auxin, n=3 embryos). Data plotted is mean +/-s.e.m. **c**, Cumulative percentage of nuclei exhibiting at least one PP7 burst by the indicated time. Conditions and data plotted as in **b. d**, Distribution of transcriptional burst sizes in the indicated conditions (embryos from **b, c**). Boxplots indicate the median (white line), interquartile range (IQR; black box), and 1.5 x IQR (whiskers). P-values were calculated using a two-sided Mann-Whitney U-test; n indicates the total number of bursts analyzed. **e**, Percentage of nuclei (% of simulations), from polymer simulations, with at least one burst at the indicated time (simulation step) in the indicated conditions. Data show is mean and 95% C.I. **f**, Loop strength ratios (ΔNIPBL/WT) of a common list of loops (**Supplementary Table 1**) in merged micro-C datasets, binned by loop length. ‘n’ is the number of loops in each bin. Box plot shows the median and IQR, whiskers extend from 10^th^-90^th^ percentiles. Distributions were compared to a null distribution (mean=1) using two-sided Wilcoxon signed-rank tests. P-values were 0.08, 0.08 and 0.02 (significance indicated by **\***) for 0-30 kb, 30-70 kb and 70-250 kb bins respectively. **g**, Loop strength ratio (as in **f**) for the 240 kb *scyl-chrb* tethering loop. Each dot represents the ratio from a biological replicate. Bar shows the median.

Polymer simulations without the sticky tether monomers also resulted in less persistent E-P contacts, resulting in reduced numbers of activated nuclei (**Fig. 3e**). As loop extrusion and adhesion facilitate different steps in E-P communication in this model (search and contact-residence, respectively), removal of loop extrusion and the distal tether resulted in a nearly complete loss of *chrb* transcription (**Fig. 3e**). Removal of the tether combined with depletion of NIPBL resulted in a similar loss of *chrb* expression (**Fig. 3b, 3c, Supplementary Fig. 4a**). This model also shows that stronger tethers do not exert much of an effect on gene expression in the absence of loop extrusion (**Supplementary Fig. 4e**).

The simultaneous loss of loop extrusion and the distal tethering element resulted in a substantial reduction in burst size (**Fig. 3d**), and this reduction was recapitulated in our model (**Supplementary Fig. 4f**). This was due to shorter estimated ON durations (**Supplementary Fig. 4c**) and was accompanied by longer estimated OFF durations (**Supplementary Fig. 4d**). These observations suggest that either cohesin or tethers are sufficient to ensure essentially normal burst durations. Loss of E-P contact stability is strongly correlated with the *scyl-chrb* distance, as removing both the distal tether and nearby CTCF-binding site resulted in a strong increase in E-P distances as measured by quantitative DNA FISH assays (**Supplementary Fig. 4g**).

Our model suggests that loop extrusion enables more frequent tether-tether contacts by reducing their effective distance. We therefore performed micro-C assays on NIPBL depleted embryos to examine tether-tether contacts across the *Drosophila* genome. Indeed, we observed a preferential weakening of long-distance interactions (**Figure 3f, Supplementary Fig. 5a**), including a 50% reduction in *scyl-chrb* focal contacts (**Figure 3g, Supplementary Fig. 5b**). Moreover, long-range contacts located near a CTCF ChIP-seq peak display increased sensitivities to NIPBL depletion (**Supplementary Fig. 5c, 5d**).

These observations are consistent with the role of NIPBL in long-range E-P interactions in cultured mammalian cells. However, in contrast to mammalian systems, NIPBL depletion resulted in very few overall changes in TADs or compartmentalization (**Supplementary Fig. 5e-g**), which is consistent with previous studies using cultured Kc167 cells^57,58^. It would therefore appear that NIPBL/cohesin is important for specific long-range associations in *Drosophila*, but not global 3D genome organization. In mammals, both processes depend on cohesin, making it difficult to tease apart the direct contribution of loop extrusion to long-range E-P contacts.

### Longer cohesin loops can compensate for the loss of tethers

These data strongly suggest that both the search time for E-P encounters and the stability of encounters are important for long range E-P communication. We next asked what perturbations might improve E-P communication and increase the speed and/or number of cells that respond. While stronger tether interactions might be expected to improve bursting probabilities, simulated changes in tether strength only had modest effects on transcription due to infrequent contacts in the absence of loop extrusion (**Supplementary Fig. 4e**). By contrast, increasing the length of cohesin loops resulted in a substantial increase in the fraction of expressing cells due to greater proximity of tethering elements (**Supplementary Fig. 3f**).

We tested these predictions experimentally in living embryos. Prior studies in mammalian cells have shown that depletion of WAPL results in less cohesin unloading and longer loops^6,59^. We reduced WAPL levels using heterozygous embryos with one functional allele of *wapl* (*wapl*^*2*^ /+, hereafter *wapl*^*2*^). *wapl*^*2*^ micro-C contact maps showed a substantial increase in contacts within ~200 kb, consistent with the formation of longer cohesin-mediated loops (**Fig. 4a; Supplementary Fig. 6a**) without changing compartmentalization (**Supplementary Fig. 6b**). Moreover, ~16% of boundaries showed a greater than 4-fold change in insulation scores (**Supplementary Fig. 6c-e**). WAPL heterozygotes also exhibited changes in loop strength, whereby shorter loops (<30 kb) are reduced in loop strength while longer loops were strengthened (**Fig. 4b**). We also observed a few new loops at novel TAD boundaries (**Supplementary Fig. 6f**).

**Fig. 4.**
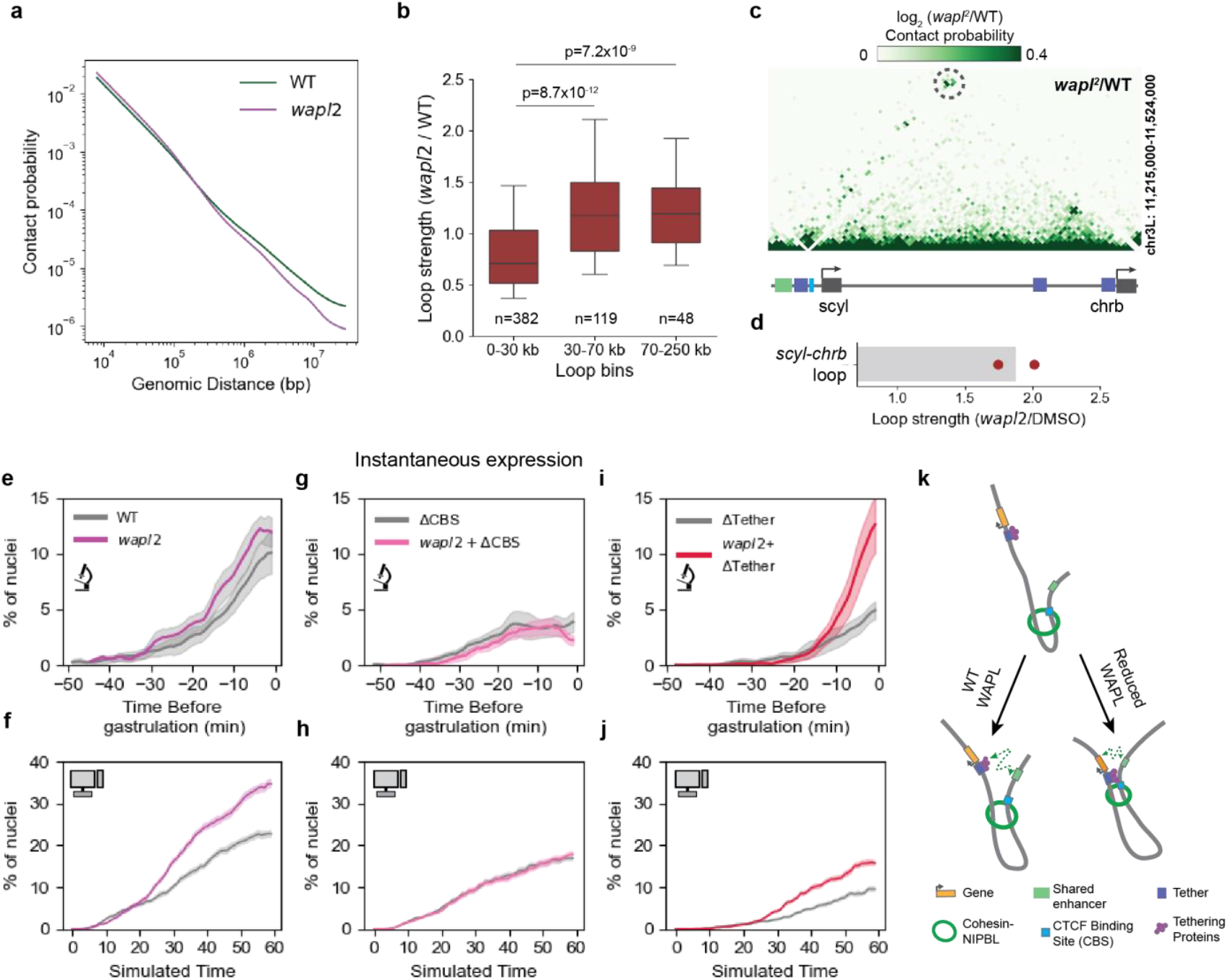
Longer cohesin loops can increase E-P contacts in the absence of tethers. **a**, Contact probability decay profiles for chr3L in WT (Tn5 library-2^nd^ replicate) and *wapl*^*2*^*/FM7c* (merged) NC14 micro-C contact maps (1.6 kb resolution). **b**, Loop strength ratios (*wapl*^*2*^/WT) for a common loop set (**Supplementary Table 1**) binned by loop length. ‘n’ is the number of loops in each bin. Box plot shows the median and IQR, whiskers extend from 10^th^-90^th^ percentiles. P-values were obtained by comparing the distributions using a two-sided Mann-Whitney U-test against the 0-30 kb bin. **c**, Differential plot (*wapl*^*2*^/WT) of micro-C maps (at 1.6 kb resolution) at the *scyl/chrb* locus showing the increase in tether-tether contacts. The *scyl-chrb* loop is circled. **d**, Loop strength ratios (as in **b**) for the 240 kb *scyl-chrb* loop. Each dot is from a biological replicate. Bar shows the median. **e, g, I**, Percentage of nuclei with a PP7 burst at the indicated time for WT (same as **Fig. 1f**), *wapl*^*2*^ (**e-** *wapl*^*2*^/+ with WT *scyl/chrb* allele, n=8), ΔCBS (same as **Fig. 2b**) and *wapl*^*2*^ + ΔCBS (**G-** *wapl*^*2*^/+ with ΔCBS *scyl/chrb* allele, n=7), ΔTether (same as **Fig. 3c**) and *wapl*^*2*^ + ΔTether (**i-** *wapl*^*2*^/+ with ΔTether *scyl/chrb* allele, n=8), Data shown is mean +/-s.e.m. **f, h, j**, Polymer simulation results showing the percentage of nuclei (% of simulations), corresponding to the conditions in the panels above. Data show is mean and 95% C.I. **k**, Cartoon model showing reduced E-P distance due to longer cohesin loops in wapl heterozygotes.

As expected from our model, reduced levels of WAPL increased micro-C contacts between *scyl* and *chrb* (**Fig. 4c, 4d**). We therefore measured *chrb* transcription dynamics in embryos derived from *wapl*^*2*^*/+* females. There is a slight increase in expression of the WT allele (*wapl*^*2*^) (**Fig. 4e, Supplementary Fig. 7a**) due to the predicted 2-fold increase in cohesin stability and a mean loop size of ~200 kb (**Fig. 4f, Supplementary Fig. 7d**). Augmented *chrb* transcription depends on the enhancer-adjacent CTCF-anchor (*wapl*^*2*^ + ΔCBS **Fig. 4g, Supplementary Fig. 7b**) as predicted by the model (**Fig. 4h, Supplementary Fig. 7e**). Reduced WAPL levels also showed no changes in the modeled ON and OFF durations for the wildtype and ΔCBS alleles (**Supplementary Fig. 7d, 7e**), confirming that the changes in transcription are a result of more frequent E-P contacts but not more stable E-P loops.

The increased frequency of E-P contacts implied by the micro-C maps and our transcription data raised the possibility that loss of the distal tether could be compensated by a reduction in WAPL. Strikingly, we observed rescue of expression in the ΔTether allele upon reduction of WAPL (*wapl*^*2*^ + ΔTether). Instantaneous transcription of *chrb* nearly reached wild-type levels in the double mutant (**Figure 4i, Supplementary Fig. 7c**) as predicted by the model (**Fig. 4j, Supplementary Fig. 7f**). The *wapl*^*2*^ + ΔTether embryos showed no significant increase in ON-durations and somewhat reduced OFF-durations compared to ΔTether alone (**Supplementary Fig. 7g, 7h**), consistent with a faster search time due to increased E-P proximity in the presence of longer loops (**Fig. 4k**).

## DISCUSSION

The role of loop extrusion in genome folding has been well characterized in many systems. Here we explored the role of this process in specific long-range interactions within TADs. We combined live transcription imaging, genetic manipulations of NIPBL and WAPL, and polymer modeling to show that loop extrusion cooperates with ‘sticky’ elements (e.g., tethers) to increase the frequency of E-P associations. These observations suggest that tuning loop extrusion and the stickiness of enhancer-proximal and promoter-proximal elements can be used to regulate the dynamics of gene expression.

The absence of loop extrusion signatures in the Hi-C maps of invertebrates has raised doubts about the occurrence of loop extrusion processes in these organisms, even though *Nipped-b* gene was identified in a genetic screen for regulators of long-distance enhancer activity in Drosophila^60^. Here, we have provided strong evidence that *Drosophila* NIPBL is essential for long-distance enhancer activity in a manner consistent with previous reports in vertebrates (**Fig. 1, Supplementary Fig. 8**). Furthermore, the role of the positioned CTCF anchor (**Fig. 2**) and the impact of reduced levels of WAPL (**Fig. 4**) are all consistent with the proposed roles of these factors as regulators of loop-extrusion in vertebrates. The differences in *Drosophila* boundary proteins and the high density of transcribing genes and insulators may explain the different roles of loop extrusion in the formation of TADs and TAD boundaries in vertebrates and flies^24,50,61,62^.

Increased gene expression is often, but not always, associated with increased E-P proximity^12,13,63,64^. The nature of E-P contacts and the role of cohesin in these contacts persist as mysteries due to the lack of direct observations of transcription dynamics. We have shown that there are no significant changes in ON-durations or burst sizes upon NIPBL-depletion (**Fig. 2e, Supplementary Fig. 2g**) or in WAPL heterozygotes (**Supplementary Fig. 7g**). By contrast, there are substantial changes in the number of nuclei exhibiting expression (**Fig. 1f, 4i**). These observations strongly suggest that cohesin enables E-P searches but is not required for the stability of E-P contacts once formed. This view is consistent with recent single-cell live transcriptional imaging in mouse ES cells and RNA-FISH assays in macrophages^20,65^.

Although loop extrusion was thought to be the primary mechanism for long-distance enhancer-mediated gene regulation in mammalian systems, essential genes such as *Sox2* and *Shh* possess additional cis-regulatory elements to ensure timely E-P contacts^9,66–68^. In vertebrates there is emerging evidence that long-distance enhancers are sensitive to depletion of potential looping proteins such LDB1^14,69^. In flies tethering elements can enable efficient E-P contacts in the absence of loop extrusion (**Fig. 3**).

In summary, we propose a “scan and snag” model for long-range gene control whereby loop extrusion primarily determines the frequency of E-P contacts and increases the likelihood of interactions between “sticky” looping factors bound at enhancers and their target promoters. A desirable feature of this model is the ability to tune E-P interactions in a tissue-specific manner through the use of different members of families of looping factors, such as POZ-domain protein families in flies and YY1, LDB1 and LHX protein families in mammals.

## Acknowledgments

We thank all members of the Levine and Boettiger labs, especially A. Mariossi and X. Li, for helpful discussions. We thank J. Raimundo for the MS2 and PP7 tagged fly lines. The work was supported by National Institutes of Health grant R35 GM118147 (to M.S.L.), NIH award U01DK127419 (to A.N.B) and NSF grant EF2022182 (to A.N.B.).

## Data Availability

The micro-C data generated in this study was uploaded to GEO (Accession number-GSE318118 to be made public upon journal publication). Published chromatin immunoprecipitation-sequencing (ChIP-seq) data for CTCF, GAF and Cp190 is available at EBI (Accession number E-MTAB-9156). CTCF CUT & TAG data is available at EBI (Accession number E-MTAB-11993). *yw* micro-C data was generated by merging NC14 datasets from GEO (GSE173518 and GSE171396).

## Code Availability

The custom pipelines for image analysis and micro-C alignment and processing can be found on github (https://github.com/pavancss/LongRangeGeneRegulation). Code for the polymer simulations can be found at this link https://github.com/tee-udon/boettiger_livecell. Brief descriptions can be found in the methods section. Any additional code is available upon request.

---

**Supplementary Fig. 1.**
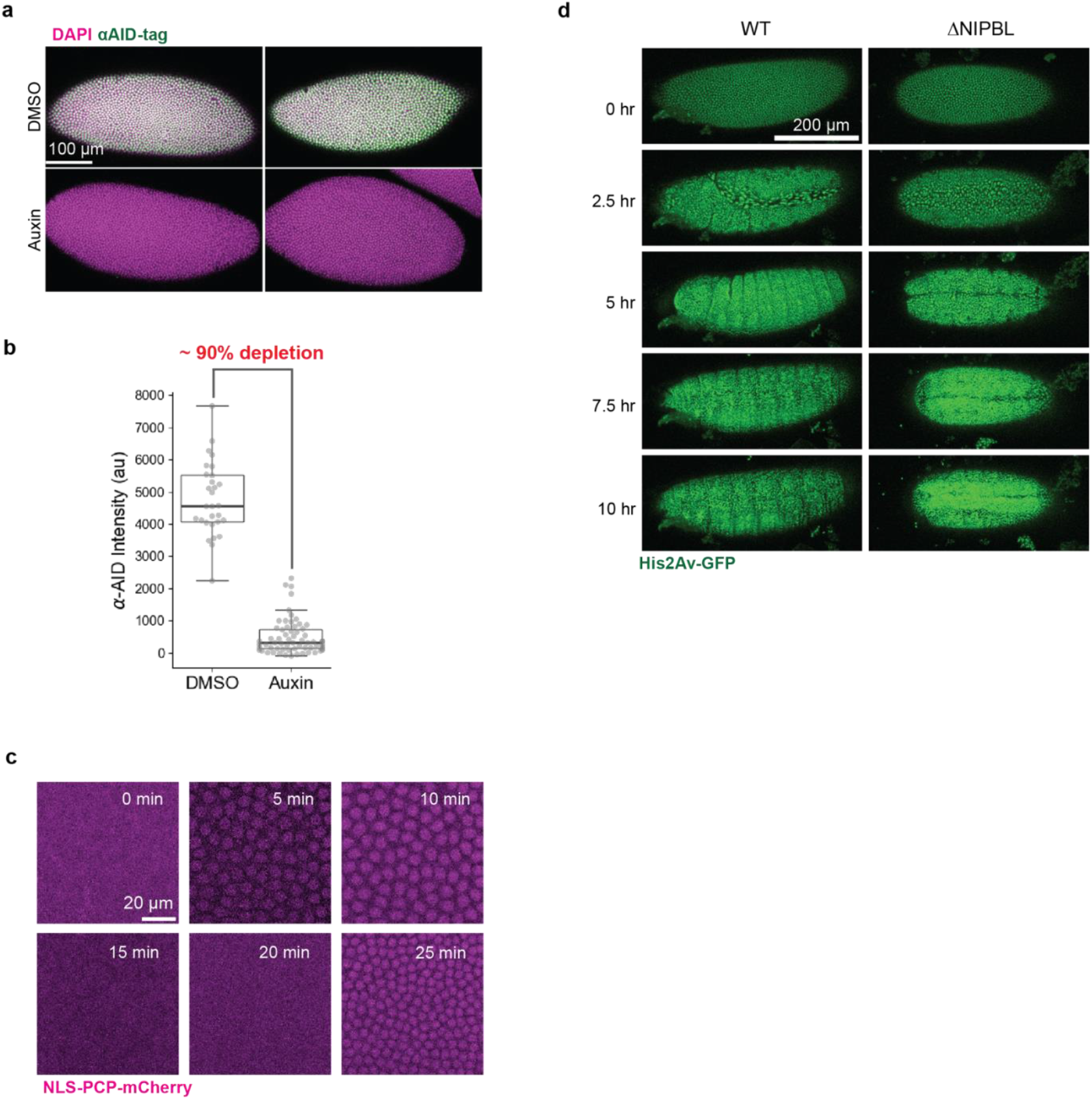
Characterization of the NIPBL depleted embryos. **a**, Representative immunofluorescence images of *NIPBL-AID-mCherry*;*nanos>OsTIR(F74G)* embryos treated with DMSO or auxin. Tagged NIPBL was detected using AID-tag antibody. **b**, Quantification of depletion efficiency showing ~90% reduction in AID-tag signal. Each dot represents the background-corrected mean intensity of a single embryo (n=29 DMSO and n=63 auxin). **c**, Representative images of NIPBL-depleted embryos undergoing two mitotic divisions, visualized using NLS-PCP-mCherry. **d**, Time-lapse imaging of *NIPBL-AID-mCherry*; *His2Av-GFP* / *nanos>OsTIR1(F74G)* embryos treated with DMSO (WT) or 50 μM auxin (ΔNIPBL). NIPBL-depleted embryos exhibit no obvious mitotic defects but fail to hatch.

**Supplementary Fig. 2.**
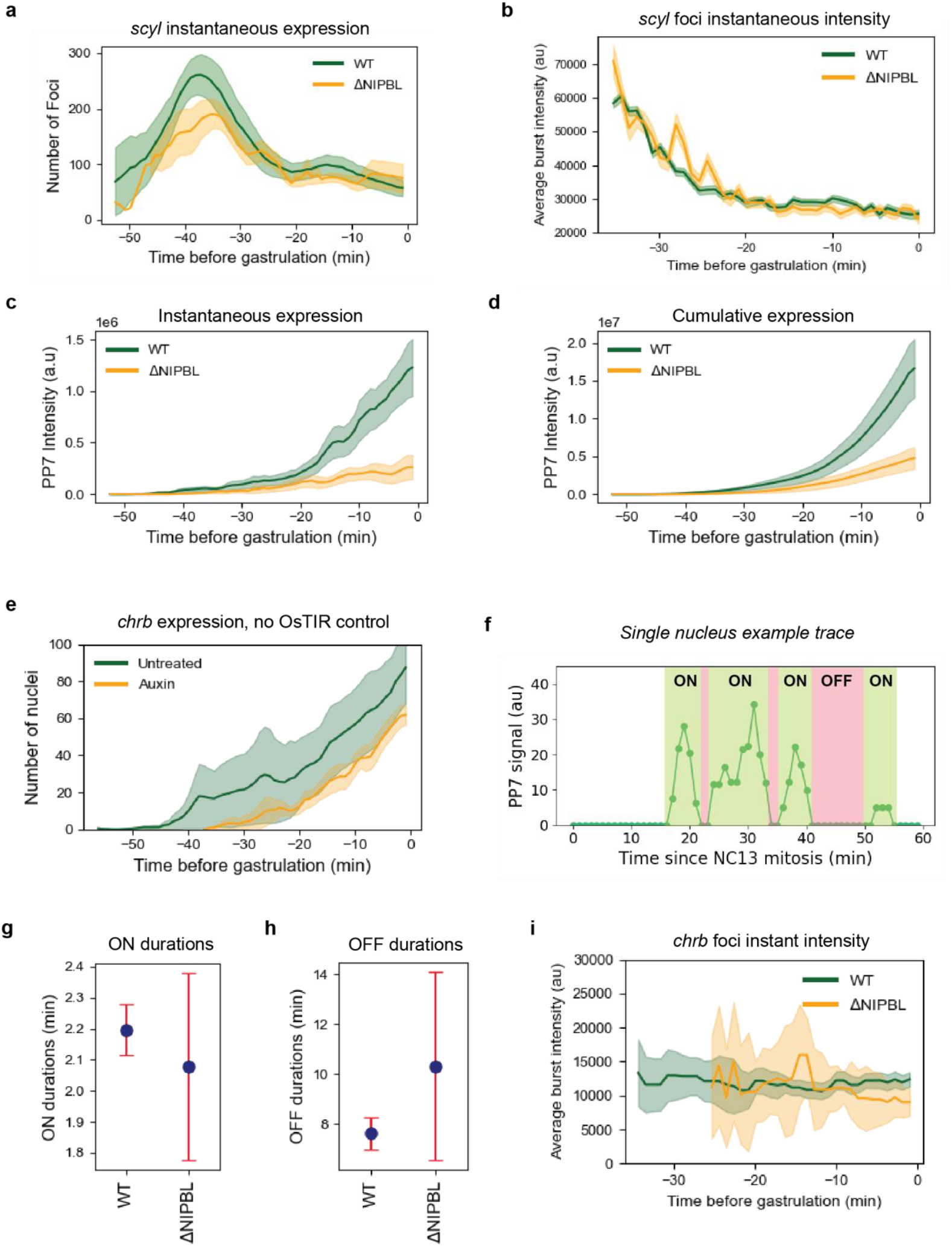
NIPBL depletion affects long-range enhancer activity. **a**, Number of MS2 foci (*scyl*) over time, showing no significant effect of NIPBL depletion on *scyl* expression. **b**, Instantaneous intensity of individual MS2 foci (*scyl*) in WT and ΔNIPBL embryos. Data plotted is mean +/-s.e.m for ≥ 3 embryos at each time point. **c**, Sum of intensities of all the PP7 foci (*chrb*) in an embryo at the indicated time for WT and ΔNIPBL embryos. Data is from the same embryos as **Fig. 1e**. Data plotted is mean +/-s.e.m. **d**, Cumulative PP7 expression (*chrb*) in WT and ΔNIPBL embryos. Data plotted is from the same embryos as **Fig. 1e**. Data plotted is mean +/-s.e.m. **e**, Number of nuclei showing PP7 expression in control embryos (no OsTIR expression) treated with DMSO (n=4) or Auxin (n=3). Data plotted is mean +/-s.e.m. **f**, Schematic defining transcriptional “ON” and “OFF” periods in a representative single-nucleus PP7 trace. **g**, Modeled ON durations in WT (n=2211 bursts) and ΔNIPBL (n=317 bursts) embryos. Data plotted is mean and error bars are 95% C.I. **h**, Modeled OFF durations in WT (n=1769 pauses) and ΔNIPBL (n=275 pauses) embryos. Data plotted is mean and error bars are 95% C.I. **i**, Instantaneous intensity of individual PP7 foci (*chrb*) in WT and ΔNIPBL embryos. Data plotted is mean +/-s.e.m for ≥ 10 bursts at each time point.

**Supplementary Fig. 3.**
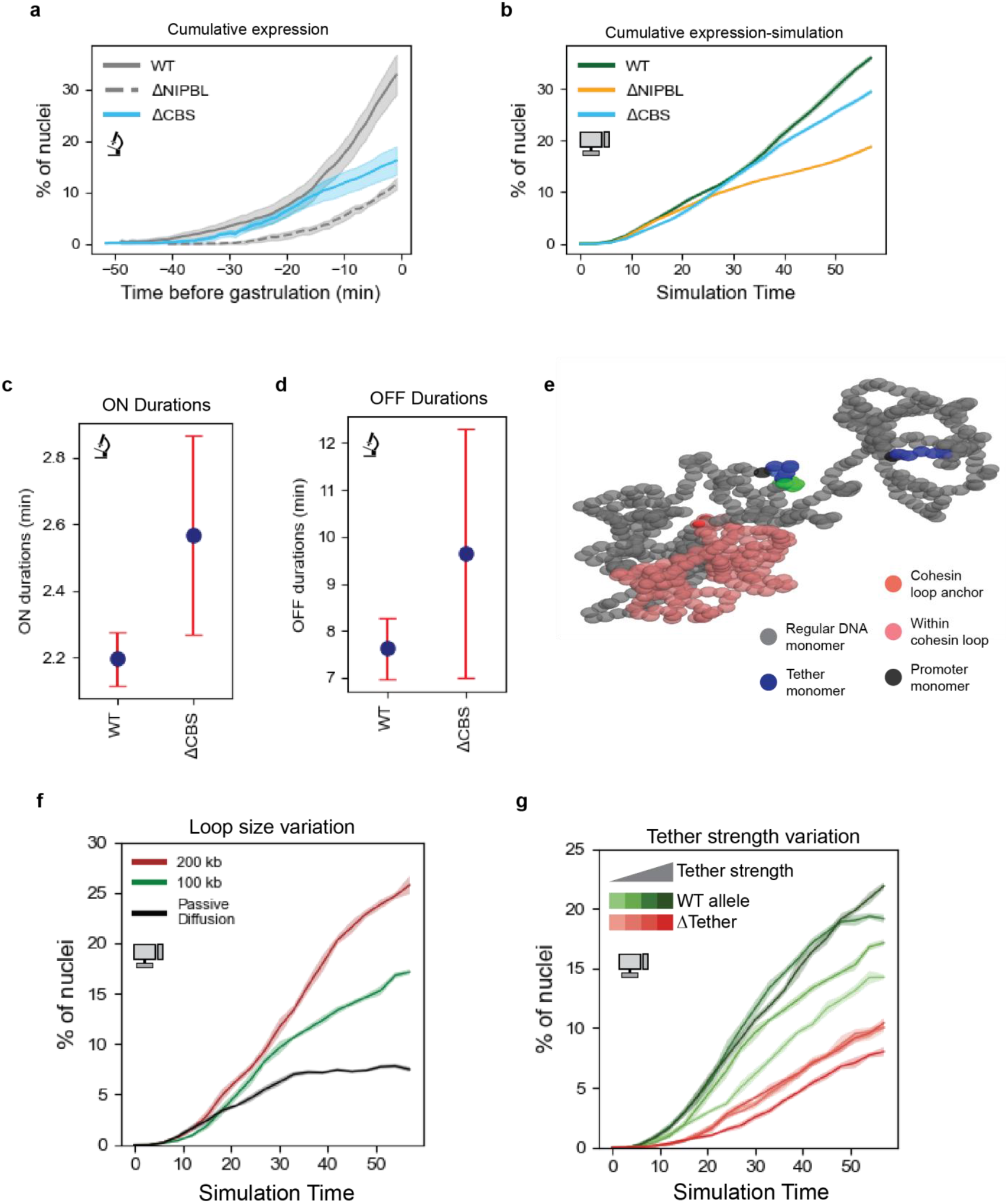
CTCF-anchored loop extrusion enables long-range gene regulation. **a**, Cumulative percentage of nuclei with PP7 foci (*chrb*) by the indicated time. The embryos analyzed are the same as shown in **Fig. 2b**. Data shown is mean +/-s.e.m. **b**, Cumulative percentage of nuclei (% of simulations), from polymer simulations, with at least one burst by the indicated time (simulation step) in the indicated conditions. Data show is mean +/-s.d. **c**, Modeled ON-durations in the indicated conditions. WT data is as shown in **Supplementary Fig. 2g**. Number of bursts analyzed was 450 in ΔCBS embryos. Marker shows the mean; error bars are 95% C.I. **d**, Modeled OFF-durations in the indicated conditions. WT data is as shown in **Supplementary Fig. 2h**. 583 pauses were analyzed in ΔCBS embryos. Marker shows the mean; error bars are 95% C.I. **e**, Snapshot of polymer model showing the effect of loop extrusion on E-P contacts. **f**, Percentage of simulated nuclei with expression over time with a fixed tether strength (35 monomers). Passive diffusion (ΔNIPBL) was compared with models with mean loop sizes of 100 kb and 200 kb. Data plotted is mean +/-s.d. **g**, Percentage of simulated nuclei with expression over time with varying tether strengths (23, 30, 35 and 40 monomers) in fibers with (WT allele) or without distal tether (ΔTether).

**Supplementary Fig. 4.**
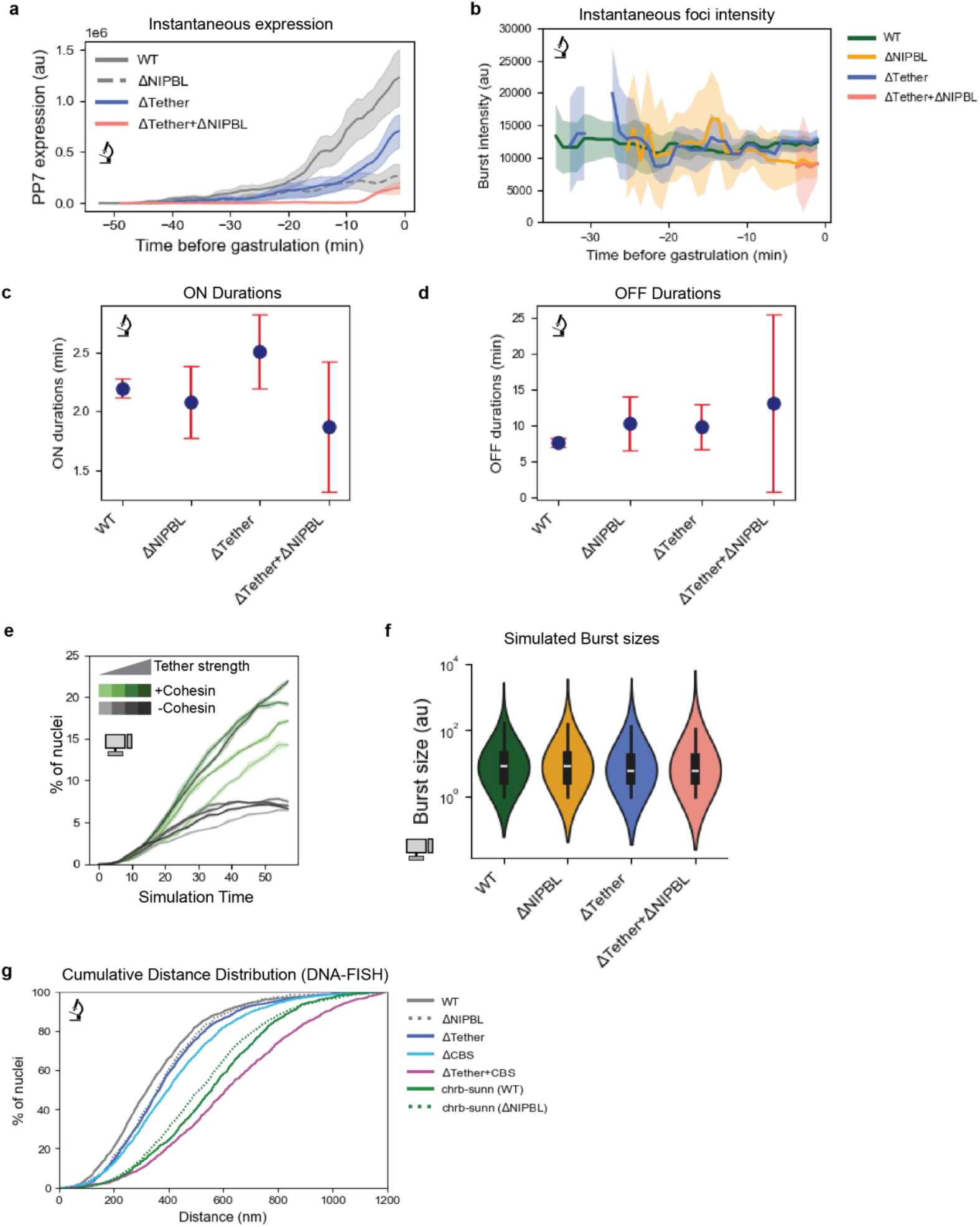
Cohesin and tethers play different roles in long-range E-P contacts. **a**, Sum of intensities of all the PP7 foci (*chrb*) in an embryo at the indicated time for WT and ΔNIPBL embryos. Data is from the same embryos as **Fig. 3b**. Data plotted is mean +/-s.e.m. **b**, Instantaneous intensity of individual PP7 foci (*chrb*) in the indicated conditions. Data plotted is mean and s.e.m for a minimum of 10 bursts at each time point. **c**, Modeled ON-durations in the indicated conditions. WT and ΔNIPBL data is as shown in **Supplementary Fig. 2g**. Number of bursts analyzed was 470 for ΔTether and 65 for ΔTether+ΔNIPBL. Point shows the mean and the error bars are 95% C.I. **d**, Modeled OFF-durations in the indicated conditions. WT and ΔNIPBL data is as shown in **Supplementary Fig. 2h**. Number of pauses analyzed was 411 for ΔTether and 42 for ΔTether+ΔNIPBL. Point shows the mean and the error bars are 95% C.I. **e**, Simulation data showing the percentage of nuclei with expression in polymers with varying tether strengths (23, 30, 35 and 40 monomers) with 100 kb cohesin loops and passive diffusion only (-Cohesin). **f**, Transcriptional burst sizes for the polymer simulations in **Fig. 3e**. Conditions and data plotted as shown. **g**, Cumulative distribution of the 3D-distance between *scyl-chrb* and *chrb-sunn* in the indicated conditions. The number of nuclei analyzed, the change in median distance and the p-values vs WT are as follows: ΔTether (1835 nuclei, 36.7 nm, p-value=2.4e-6), ΔCBS (1700 nuclei, 71 nm, p-value=3.8e-18) and ΔTether+ΔCBS (1298 nuclei, 287 nm, p-value=1.38e-145).

**Supplementary Fig. 5.**
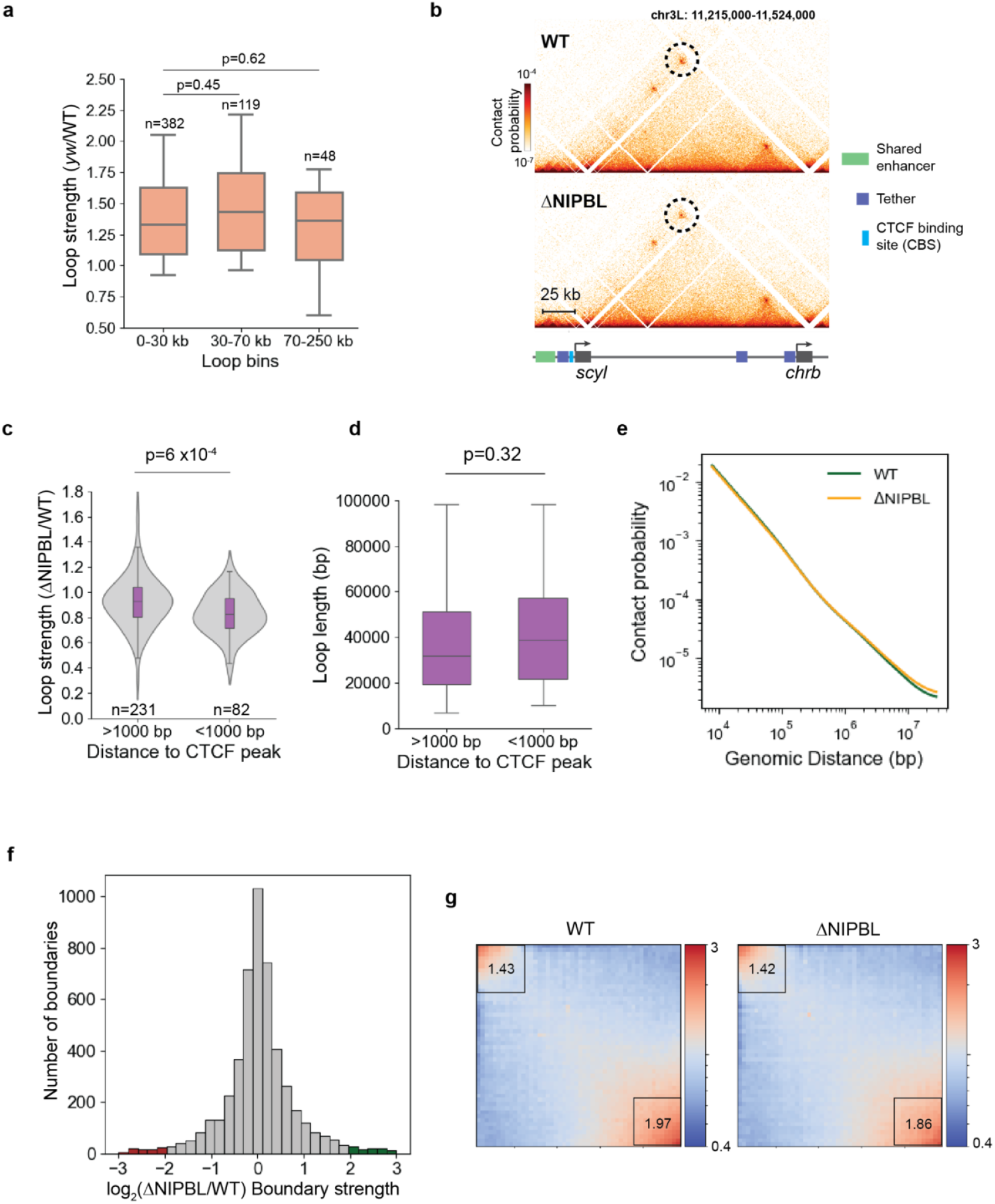
Loop extrusion enables long-distancer tether-tether interactions but is dispensable for overall genome organization in fly embryos. **a**, Loop strength ratios (*yw*/WT) of a common list of loops (**Supplementary Table 1**) in merged micro-C datasets, binned by loop length. ‘n’ is the number of loops in each bin. Box plot shows the median and IQR, whiskers extend from 10^th^-90^th^ percentiles. Distributions were compared using a two-sided Mann-Whitney U-test and the p-values are shown in the plot. **b**, Micro-C maps (800 bp resolution) of the *scyl-chrb* locus in merged WT (DMSO) and ΔNIPBL (Auxin) NC14 embryos. The *scyl-chrb* loop is circled. **c**, Loop strength ratios of a common list of loops (**Supplementary Table 1**) in merged WT and ΔNIPBL micro-C binned by distance of anchors to a CTCF ChIP-seq peak. ‘n’ is the number of loops in each bin. Box plot shows the median and IQR. The whiskers extended from 10^th^-90^th^ percentiles. Distributions were using a two-sided Mann-Whitney U-test and the p-value is shown in the plot. **d**, Loop lengths in the two bins in **c**. The box shows the median and interquartile range (IQR-25-75^th^ percentile) and the whiskers are plotted at 1.5 IQR. The median of the two distributions was compared using a two-sided Mann-Whitney U-test and the p-value is shown. **e**, Contact probability decay profiles for chr3L in WT and ΔNIPBL (2^nd^ replicate) NC14 micro-C contact maps (1.6 kb resolution). **f**, Histogram of the boundary strength ratio of strongly insulating loci showing very few changes in insulation strength. Boundaries with >4-fold change in boundary strength are shaded (red-reduced, green-increased). **g**, Saddle plots showing compartmentalization in WT and ΔNIPBL micro-C maps. The interaction strength between active regions (AA-lower right box) and inactive regions (BB-upper left box) are shown.

**Supplementary Fig. 6.**
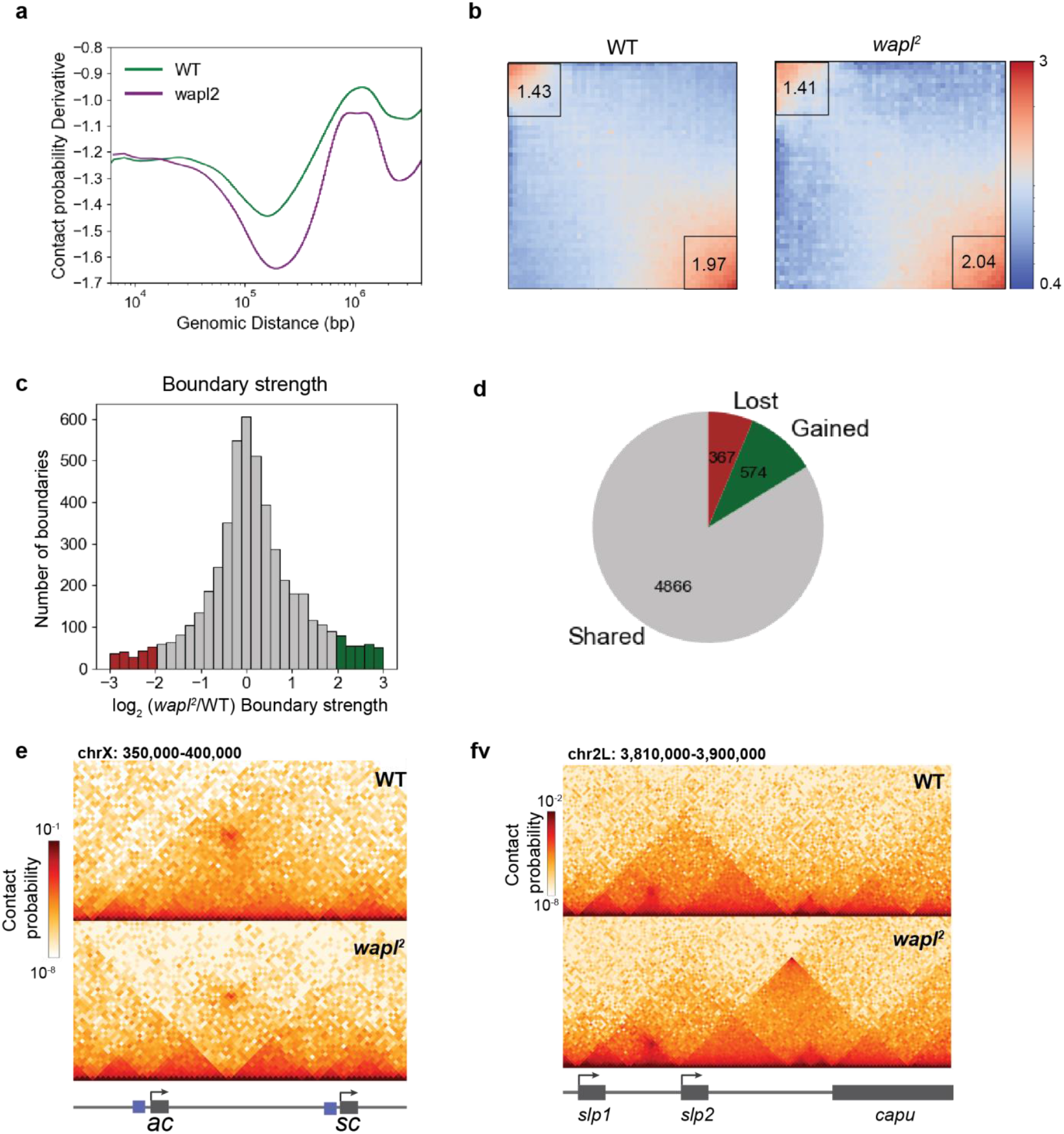
Reduced WAPL levels show significant changes in genome organization. **a**, Differential contact probability curves in WT and *wapl*^*2*^ micro-Cmaps. **b**, Saddle plot showing weak and unchanged compartmentalization in WT and *wapl*^*2*^ micro-C maps. The interaction strength between active regions (AA-lower right box) and inactive regions (BB-upper left box) are shown. **c**, Histogram of the boundary strength ratio of strongly insulating loci showing changes in insulation strength. Boundaries with >4-fold change in boundary strength are shown in color (red-reduced in *wapl*^*2*^ maps, green-increased in *wapl*^*2*^ maps). **d**, Pie-chart showing number of shared (grey), lost (>4-fold reduction in *wapl*^*2*^ maps) and gained (>4-fold increase in *wapl*^*2*^ maps) boundaries. **e**, Micro-C maps of the *ac-sc* locus showing a new boundary in the *wapl*^*2*^ micro-C map. **f**, Micro-C maps of the *slp1-slp2* locus showing a new TAD and loop formed between the TAD boundaries in the *wapl*^*2*^ sample.

**Supplementary Fig. 7.**
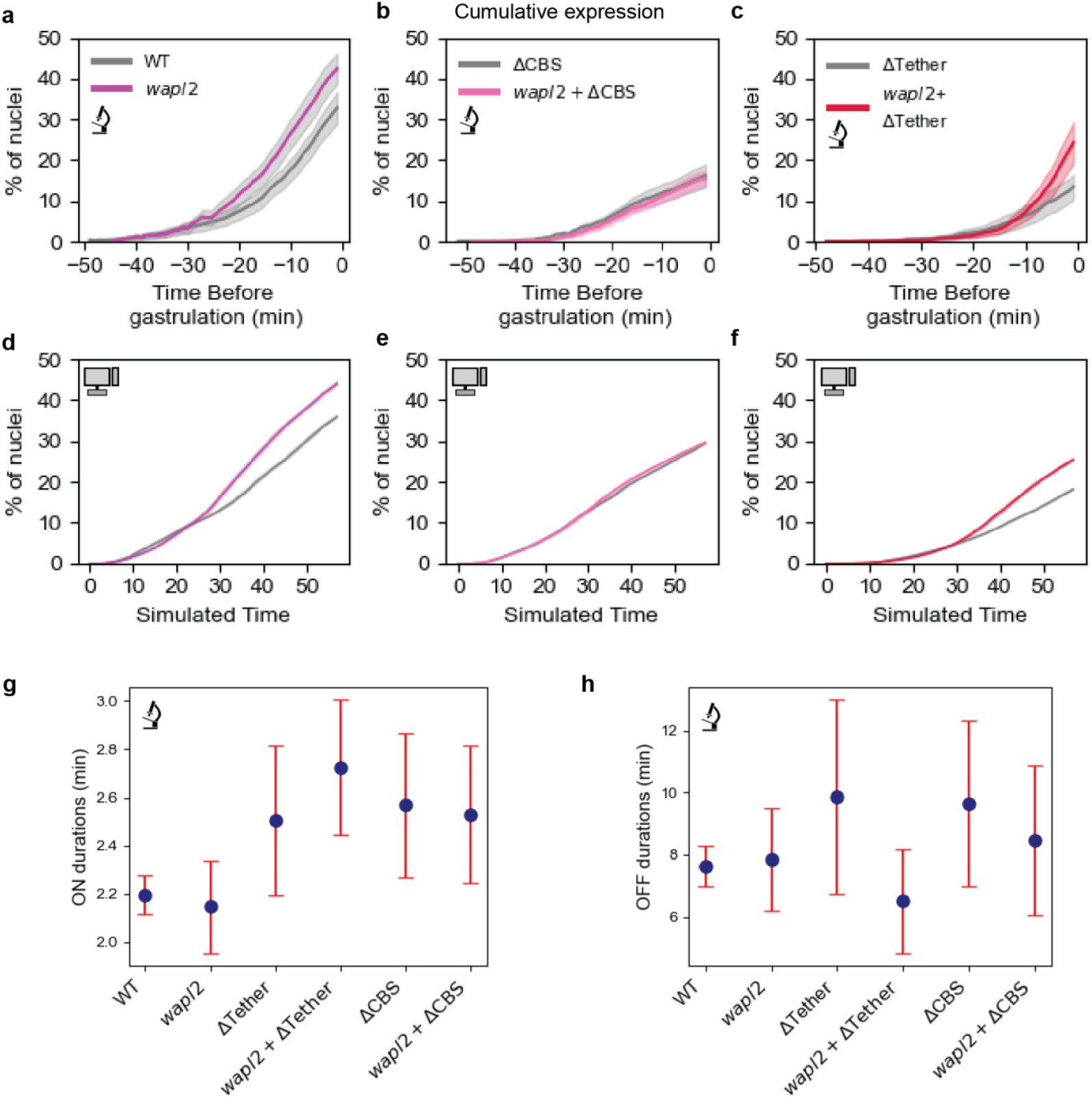
Reduced WAPL levels increase E-P contact frequency. **a, b, c**, Cumulative percentage of nuclei with a PP7 burst by the indicated time for WT (**a-** WT allele + DMSO, same as in **Fig. 1e**), *wapl*^*2*^ (**a-** same embryos as **Fig. 4e**), ΔCBS (**b-** same as **Supplementary Fig. 3a**) and *wapl*^*2*^ + ΔCBS (**b-** same embryos as **Fig. 1f**), ΔTether (**c-** same as **Fig. 3b**) and *wapl*^*2*^ + ΔTether (**c-** same embryos as **Fig. 4i**), Data shown is mean +/-s.e.m. **d, e, f**, Polymer simulation results showing cumulative expression corresponding to the conditions in the panels above. Data show is mean +/-s.d. **g**, Modeled ON-durations in the indicated conditions. WT and ΔCBS and ΔTether data is as shown in **Supplementary Fig. 2g, 3b** and **4c** respectively. Number of bursts analyzed was 2034 for *wapl*^*2*^, 572 for *wapl*^*2*^ + ΔCBS and 801 for *wapl*^*2*^ + ΔTether. Point shows the mean and the error bars are 95% C.I. **h**, Modeled OFF-durations in the indicated conditions. WT and ΔCBS and ΔTether data is as shown in **Supplementary Fig. 2h, 3c** and **4d** respectively. Number of pauses analyzed was 2058 for *wapl*^*2*^, 575 for *wapl*^*2*^ + ΔCBS and 801 for *wapl*^*2*^ + ΔTether.

**Supplementary Fig. 8.**
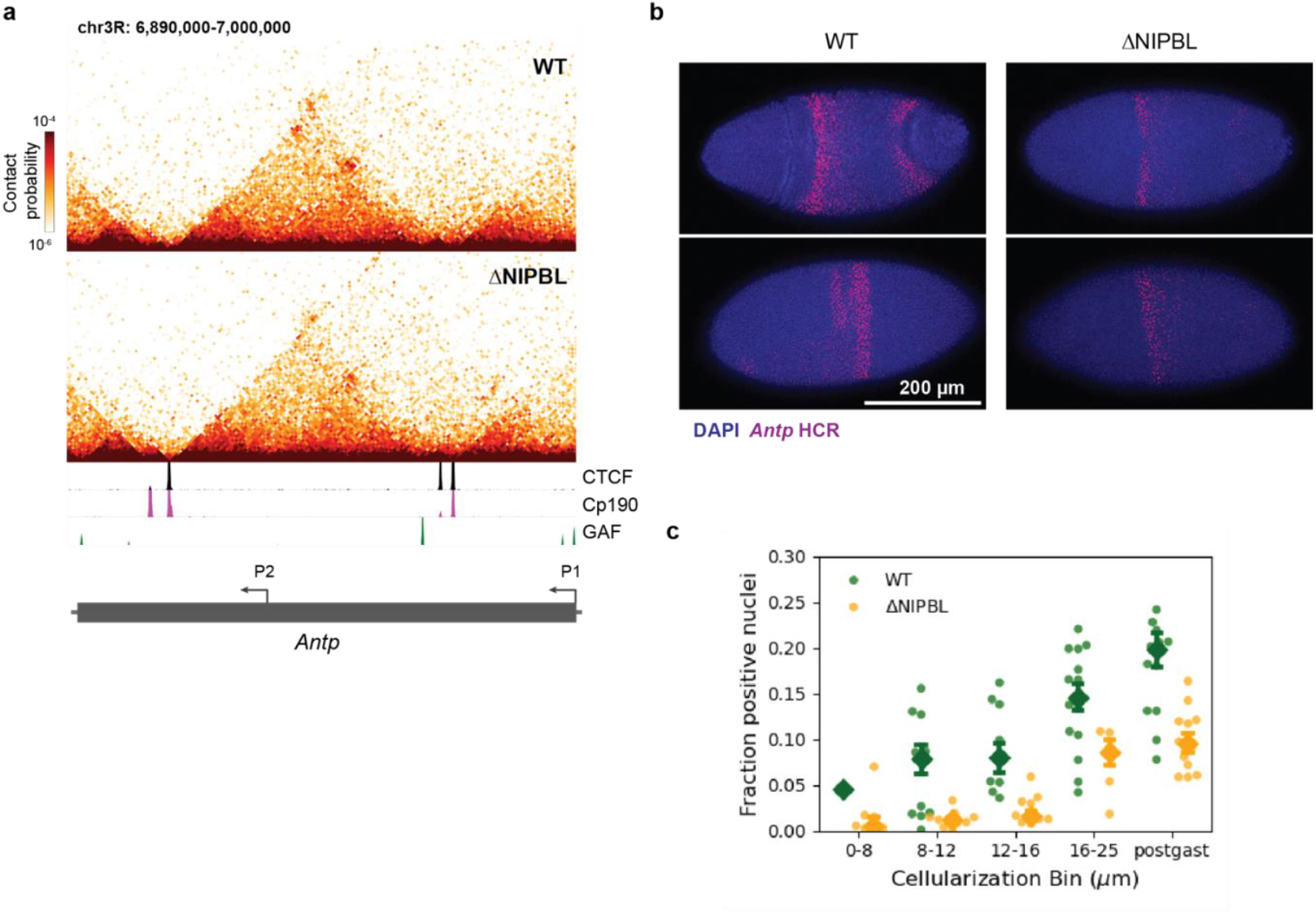
NIPBL depletion weakens loops and expression at *Antp* locus. **a**, Micro-C maps showing the change in loop strength at the *Antp* locus. ChIP-seq tracks of CTCF, Cp190 and GAGA-associated factor (GAF) are from nuclear cycle 14 (NC14) *Drosophila* embryos^61^. **b**, *In situ* hybridization chain reaction (HCR) for *Antp* RNA (common intron shared between P1 and P2 promoters) in WT and ΔNIPBL embryos. *Antp* expression in the early embryo is driven by enhancers located ~120 kb away from the P2 promoter^5^. Representative images in two orientations and similar stages are shown. **c**, Fraction of nuclei with background subtracted HCR signal in WT and ΔNIPBL embryos. The embryos were separated into bins based on embryo cellularization^70^ to account for the increase in expression of *Antp* through NC14^5^. ‘postgast’ indicates gastrulated embryos. Each dot indicates the fraction from a single embryo. The diamond marker is the median and the error bars are s.e. The expression in each bin was compared between the two conditions using a two-sided Mann-Whitney U-test and these were then combined using Fisher’s Method to assess the significance of the difference (p-value=3.46e-7).

## METHODS

### Plasmid design

#### CRISPR plasmids

The homology repair template for the AID tagging of the *NIPBL* allele was prepared using the pScarlessHD plasmid (https://www.addgene.org/64703/). An *mAID2*.*0-mCherry* cassette (https://www.addgene.org/72831/) and a *3xP3-DsRed* marker flanked by *piggybac* transposon sequences were inserted in between two 1 kb homology arms flanking the last codon of *NIPBL* (chr2R-4,737,828). The repair template was generated by ligating phosphorylated duplex guide RNA oligos, designed using flyCRISPR (http://targetfinder.flycrispr.neuro.brown.edu/index.php), into the BbsI site of pCFD3 (https://www.addgene.org/49410/).

#### OsTIR expression

The coding region of *OsTIR(F74G)* (https://www.addgene.org/140536/) was generated by replacing *MCP-GFP* coding sequence in a *nanos>MCP-GFP* plasmid^1^.

### Transgenic fly generation

#### CRISPR

MS2 and PP7 tagged *scyl* and *chrb* fly lines were generated and described previously^2^.

For endogenous NIPBL tagging, the homology repair donor and the guide RNA were injected together into a *nos-Cas9* (III) stock (BDSC #78782). Microinjections were carried out by BestGene Inc. Adults resulting from the injected eggs were crossed to *yw* flies and the progeny were screened for DsRed expression. DsRed positive single males were selected and crossed to a *gla/CyO* line to generate *CyO* balanced stocks and the individual lines were sequenced by PCR from regions flanking the homology arms to confirm a single insertion. All the studies were conducted using flies derived from a single isogenic *NIPBL-mAID-mCherry* stock.

#### Transgene insertions

*nanos>OsTIR(F74G)* plasmids were inserted into the attP sites on chr2L or chr3L by injecting the plasmid into either VK000037 (chr2L-BDSC #9752) or VK00031 (chr3L-BDSC #9748). Successful transformants were identified using *miniwhite* expression and several isogenic lines were isolated.

*NIPBL-mAID-mCherry, nanos>OsTIRF74G/CyO* flies were generated by recombining *NIPBL-mAID-mCherry* CRISPR lines with *nanos>OsTIR(F74G)* (VK37)/*CyO* flies. A single recombinant male was used to generate an isogenic stock and the *NIPBL* locus tagging was verified using PCR as described above.

*NIPBL-mAID-mCherry; nanos>OsTIR(F74G)* stock was generated similarly by recombining the *NIPBL-mAID-mCherry* CRISPR line with the *nanos>OsTIRF74G* (VK31) flies to generate a homozygous stock.

### Fly husbandry and stocks

Stocks were maintained at room temperature using standard cornmeal medium. Crosses for imaging were conducted at 25 °C on yeasted apple juice plates.

Reporter flies *NIPBL-mAID-mCherry / CyO*; nanos>*MCP-EGFP, nanos>SV40NLS-PCP-mCherry, His2Av-eBFP2 (TR)/TM6b* were generated by recombining *NIPBL-mAID-mCherry* flies with nanos>*MCP-EGFP, nanos>SV40NLS-PCP-mCherry, His2Av-eBFP2 (TR)* flies described previously^3^. The construction of *nanos>SV40NLS-PCP-mCherry, His2Av-BFP/CyO; nanos>MCP-GFP* flies used for the no OsTIR control was also reported previously^1^.

*wapl[2*^]^*/FM7c* flies were obtained from Bloomington Drosophila Stock Center (BDSC)-Stock #5741.

### Immunofluorescence

#### Fixation

Embryos were collected for 60 min from a *NIPBL-mAID-mCherry; nanos>OsTIR(F74G)* self-cross. The embryos were dechorionated in a 60 μm mesh using fresh 4.12 % sodium hypochlorite solution for 95 s. The embryos were thoroughly washed with water and then moved to a tube containing 1x DPBS supplemented with 0.25 % DMSO or 50 μM 1H-Indole-3-acetic acid, 5-phenyl-, (acetyloxy)methyl ester (5-PH-IAA-AM Cat no. HY-141894 MedChemExpress) and incubated on a nutator at 25 °C for 2 hours. The embryos were then transferred to a scintillation vial containing 5 ml of n-Heptane and 5 ml of fixative (1x DPBS + 8% formaldehyde Thermo-Scientific Cat no. 28906) and shaken vigorously at 250 rpm on a horizontal shaker for 20 min. The bottom layer containing the fixative was removed with a pipette and replaced with 10 ml of methanol and vortexed vigorously for 2-5 min to remove the vitelline membrane. The solution and any embryos that did not fall to the bottom were removed and the devitellinized embryos that collected at the bottom were washed twice with methanol. The embryos were stored in methanol at −20 °C for a few days.

#### Staining and imaging

Embryos were rehydrated using one 5 min wash with a 1:1 mix of methanol and 1xDPBS+0.1% Tween-20 (PBS-T) and then two 5 min washes with PBS-T. The embryos were then permeabilized by incubating them in 1x DPBS+0.5% Triton X-100 on a nutator for 10 min. The embryos were washed twice using PBS-T and incubated for 1 hour in Abdil (1x DPBS + 0.1% Tween-20 + 3 % bovine serum albumin) for blocking. The embryos were then incubated overnight at 4 °C with a 3 μg/ml mAID-tag antibody (ProteinTech Cat no. 28209-1-AP) or nothing (no primary control) in Abdil. The embryos were washed thrice in PBS-T the next day and then incubated with 6 μg/ml Alexa-488 conjugated Donkey anti-IgG Goat Antibody (Jackson ImmunoResearch Cat no.111-546-047) and 0.1 μg/ml DAPI for 2 hours at room temperature. The embryos were then washed thrice with PBS-T and mounted in VECTASHIELD antifade mounting medium (Vector Labs Cat no. H-1000-10). The imaging was performed using a Plan-Apochromat 20x/0.8 NA M27 objective on a Zeiss LSM 880 confocal using a two-track setup with DAPI on one track and Alexa488 on another track. 1 μm stacks were collected resulting in a voxel size of 0.52 x 0.52 μm x 0.925 μm.

### RNA-FISH

#### Probe generation

*Antp* Hybridization chain reaction (HCR) v3.0 *in situ* probes (33 pairs) were designed using Easy_HCR (https://github.com/SeuntjensLab/easy_hcr) and synthesized as an oligopool by Integrated DNA Technologies (IDT). The probes were designed against an intron downstream of *Antp* P2 promoter and reflect nascent transcription from both P1 and P2 promoters.

#### HCR protocol

HCR *in situ* hybridizations were performed, with some modifications, according to the whole mount embryo protocol provided by the manufacturer (https://files.molecularinstruments.com/MI-Protocol-HCRv3-FruitFly-Rev6.pdf)

Embryos were fixed and permeabilized as described in the immunofluorescence section. After washing with PBS-T, the embryos were treated with 4, the embryos were treated with 4 μg/ml Proteinase K for 5 min and washed thrice with PBS-T. The embryos were then incubated in 100 μl probe hybridization buffer (30% Formamide, 5x SSC, 9 mM Citric acid pH 6, 0.1 % Tween-20, 50 μg/ml Heparin, 1x Denhardts solution, 10% Dextran sulfate) for 30 min at 37 °C. 2 μl of the reconstituted probe oligopool (10 μM total oligos) was diluted into preheated 100 μl probe hybridization buffer and added to the embryos. The mixture was incubated at 37 C overnight. The buffer was removed and the embryos were washed with 200 μl probe wash buffer (30% Formamide, 5x SSC, 9 mM Citric acid pH 6, 0.1 % Tween-20, 50 μg/ml Heparin) four times at 37 °C and twice with 200 μl 5x SSC-T (0.75 M Sodium Chloride, 75 mM Trisodium Citrate Dihydrate. 0.1% Tween-20). Embryos were treated with 100 μl amplification buffer (5x SSC, 0.1% Tween-20, 10% Dextran sulfate) for 10 min while 1.5 μl each of the corresponding denatured and cooled hairpins were added to 50 μl amplification buffer and mixed with the embryos. The amplification was performed for 3 hours at room temperature. The solution was removed and the embryos washed extensively with 5x SSC-T with the final wash including 0.1 mg/ml DAPI solution. The embryos were mounted in Prolong Gold Antifade Mounting media (Invitrogen Cat no. P10144). The imaging was performed as described in the immunofluorescence section.

### Micro-C

#### Sample preparation and sequencing

##### Embryo collection and fixation

NIPBL depletion-Embryos were collected for 1 hour from either *NIPBL-mAID-mCherry, nanos>OsTIR(F74G)/CyO* x *NIPBL-mAID-mCherry* cross (First replicate) or *NIPBL-mAID-mCherry; nanos>OsTIR(F74G)* self-cross (second replicate) in a 60 μm mesh (Genessee Scientific) and dechorionated by treatment with 4.12 % Sodium Hypochlorite solution for 95 s. The embryos were washed thoroughly with deionized water and moved to a tube containing 1x DPBS (2.67 mM KCl, 1.47 mM KH_2_PO_4_, 137.93 mM NaCl, 8.06 mM Na_2_HPO_4_, Gibco Cat no. 14190-136) containing 0.25 % DMSO or 50 μM 1H-Indole-3-acetic acid, 5-phenyl-, (acetyloxy)methyl ester (5-PH-IAA-AM Cat no. HY-141894 MedChemExpress) and incubated on a nutator at 25 °C for 2 hours. The embryos were then transferred to a scintillation vial containing 5 ml of n-Heptane and 5 ml of fixative (4 ml (1x DPBS + 0.1% Triton X-100)+ 1 ml 16 % formaldehyde Thermo-Scientific Cat no. 28906) and shaken vigorously at 250 rpm on a horizontal shaker for 15 min. 3. 75 ml of 2.5 M Glycine was added to quench the fixation reaction and incubated on the shaker for 5 min. The embryos were then washed thoroughly with PBS-Tx (1x DPBS + 0.1% Triton X-100) and stored at 4 °C. Multiple batches were collected and combined for storage overnight at 4 °C. Secondary crosslinking was performed the next day by the addition of 10 ml of 1x DPBS-Tx + 3 mM Disuccinimidyl glutarate (DSG) and 3 mM ethylene glycol bis(succinimidyl succinate) (EGS) for 45 min at room temperature on a nutator. The reaction was quenched by the addition of 3.7 ml of 2M Tris-HCl pH 7.5 (Quality Biological Inc Cat no. 351048101) for 5 min. The embryos were thoroughly washed with DPBS-Tx and stored on ice for manual sorting. The embryos were hand sorted for NC14 embryos under a dissection scope and flash frozen in liquid nitrogen before storage at −80 °C.

##### Library preparation

Embryo pellets were thawed quickly on ice, resuspended in 500 μl MB1 (50 mM NaCl, 10 mM Tris pH 7.5, 5 mM MgCl_2_, 1 mM CaCl_2_, 0.2 % NP-40, 1x Protease inhibitor cocktail) and homogenized using a Dounce homogenizer. The crushed embryos were incubated on ice for 20 min, washed twice with MB1 by centrifugation and resuspended again in 500 μl MB1 buffer. 50 U of micrococcal nuclease (MNase-Worthington) was added to the embryos and the mixture was incubated at 37 °C for 10 min to digest the chromatin. The reaction was stopped by the addition of 4 mM Ethylene Glycol-bis(β-aminoethyl ether)-N,N,N’,N’-tetraacetic acid (EGTA) and incubation at 65 °C for 10 min. The nuclei were pelleted and washed twice with ice-cold MB2 (50 mM NaCl, 10 mM Tris-HCl pH 7.5, 10 mM MgCl_2_) to remove remaining MNase. The nuclei were then treated with 25 U of T4 polynucleotide kinase (New England Biolabs NEB #M0201) in end-chewing buffer (1x NEB Buffer 2.1, 2 mM ATP, 2 mM DTT) for 15 min at 37 °C and then 25 U of large Klenow fragment of DNA Polymerase I (NEB #M0210) was added and the mixture was incubated for a further 15 min at 37 °C. These DNA ends were then biotin labeled by the addition of end-labeling mixture (5 ul 1 mM Biotin-dATP, 5 ul 1 mM Biotin-dCTP, 10 mM dTTP, 10 mM dGTP, 2.5 ul 10x T4 DNA ligase buffer, 0.125 μl Bovine Serum Albumin) and incubated for 45 min at 25 °C. The labeling reaction was stopped by the addition of 30 mM EGTA and incubation at 65 °C for 20 min. The nuclei were then pelleted and washed twice with excess MB3 buffer (50 mM Tris-HCl pH 7.5, 10 mM MgCl_2_). The nuclear pellet was then incubated with 5000 units of T4 DNA Ligase (NEB #M0202) for 3 hours at room temperature to generate proximity ligated chromatin fragments. The nuclear pellet was then incubated with 500 U of Exonuclease III (NEB #M0206) to remove biotin from unligated DNA ends. Micro-C libraries for the first replicate of NIPBL depletion were then generated using the standard protocol and the libraries for the second NIPBL depletion replicates and *wapl*^2^/FM7c samples were generated using the modified protocol.

##### Standard protocol

The nuclear pellet was then incubated with 2 mg /ml Proteinase K (Invitrogen #25530049) and 1 % sodium-dodecyl sulphate (SDS) at 65 °C overnight to reverse crosslinking and digest the nuclear proteome. DNA was extracted with Phenol:Chloroform:Isoamylalcohol, ethanol-precipitated, resuspended in 50 ml of 10 mM Tris-HCl pH 8.5, treated with 2 ml RNase A at 37 °C for 30 min and purified on a ZymoClean spin column. Ligated DNA was further fragmented using sonication in a Covaris ME220 ultrasonicator to an average size of 350 bp. Biotin labeled DNA were then extracted by binding to Streptavidin beads (Dynabeads C1-Invitrogen Cat no. 65-001). The DNA ends were processed for adapter ligation by treatment with 3.5 μl of End Prep Reaction buffer and 1.5 μl End Prep enzyme mix (NEB-E7645S) in a 30 μl reaction incubate at 20 °C for 30 min and then at 65 °C for 30 min. Adapter ligation was then performed by the addition of 0.5 μl Illumina adapter, 15 μl of Ultra II Ligation Master Mix and 0.5 μl Ligation Enhancer (NEB-E7645S) and incubated for 30 min at 20 °C and then for 15 min at 37 °C following the addition of USER enzyme. The streptavidin beads were then washed in 1x BW buffer (5 mM Tris-HCl pH 7.5, 0.5 mM EDTA, 2 mM NaCl) and resuspended in 10 mM Tris-HCl pH 7.5. Library PCR was performed using KAPA HiFi HotStart Ready Mix (Roche #KK2501) and the libraries were purified using Ampure XP beads (Beckman Coulter A63880). Micro-C libraries were sequenced on a NovaSeq S1 100nt Flowcell v1.5 by the Princeton University Genomics Core Facility.

##### Modified Tn5 protocol

Nuclei were resuspended in 20 µl lysis buffer (10 mM Tris-HCl, pH 8.0; 20 mM NaCl; 1 mM EDTA; 0.1% Triton X-100; 1.5 mg/ml QIAGEN protease) and lysed by sequential incubation at 50 °C for 1 h, 65 °C for 1 h, and 70 °C for 15 min. DNA was extracted using 1.6× AMPure XP beads. The purified DNA fragments were then subjected to Tn5 tagmentation and immobilized on MyOne Streptavidin C1 Dynabeads. Following five washes, the library DNA was amplified using KAPA HiFi HotStart ReadyMix (Roche #KK2501) for 8 PCR cycles. The libraries were then sequenced using 2x 150 bp paired-end sequencing on a NovaseqX.

#### Analysis

##### Genome mapping and normalization

The sequencing data was analyzed as described previously. Briefly, the paired reads were mapped to dm6 reference genome using bwa with mem -SP5M -t8. The aligned reads, were parsed, sorted and deduplicated using pairtools with default settings. The deduplicated pairs were then selected with ‘(pair_type == “UU”) or (pair_type == “UR”) or (pair_type == “RU”)’ settings using pairtools. After being split and indexed by pairtools with default parameters, the output pairs were processed and normalized into multi-resolution matrices with default settings by Cooler.

##### Loop Strength analysis

The loop strength analysis was done on a consensus loop list that was generated from a merged embryo dataset (**Supplementary Table 1**). The consensus loop list was then filtered to remove loops present in pericentromeric regions and to remove weak loops. The remaining loops were then binned into three bins 0-30 kb, 30-70 kb and >70 kb. The loop strength was then calculated as the sum of the contacts in a 2.4 kb x 2.4 kb window and then normalized to the total number of pairs of sequences in the micro-C maps.

##### CTCF loop strength analysis

The CTCF peaks were called from CUT & TAG assay data (E-MTAB-11993). The raw data from both the wildtype (*yw*) and CTCF-depleted embryos^4^ were aligned to the reference *dm6* genome using bowtie2 (options --end-to-end --very-sensitive --no-unal --no-discordant --no-unal --no-mixed --no-discordant --phred33 -I 10 -X 700). The sorted bam files were used as treatment (*yw*) and control (CTCF-depleted) for macs3 callpeak (https://macs3-project.github.io/MACS/) command using standard options.

Loop strength was calculated as described in the previous section. The loops were binned by distance from the center of CTCF CUT & TAG peaks.

##### Contact probability, Contact probability derivative curves

Contact probability as a function of distance was obtained using the cooltools.expected_cis function from the cooltools package^5^ on 1.6 kb resolution maps. For **Supplementary Fig. 5C**, the plot was made using only chr3L contact data from the second replicate of WT and ΔNIPBL (Tn5 tagmented library). The relative contact probability decay was similar for other chromosomes and for the first replicate as well. For **Fig 4A**, the plot was made using second replicate of WT (Tn5 tagmented library) and the merged *wapl*^*2*^ map from both replicates (Tn5 tagmented libraries). Contact probability derivatives were calculated as the slope of the log(s*P(s)) vs log (s) curve for the same data as the P(s) curves.

### Live imaging

#### Auxin depletion time course

Embryos were collected from a self-cross of *NIPBL-mAID-mCherry*; *nanos>OsTIR(F74G)* flies for 2 hours, dechorionated for 100 s with fresh 4.12 % sodium hypochlorite solution and washed thoroughly with water. Excess water was wicked out and the embryos were transferred onto a 35 mm glass bottom dish (Mattek Cat no. P35G-1.5-20-C). 3 ml of 1x DPBS containing 50 μM 5-PH-IAA-AM was added gently to the dish to prevent dislodging the embryos from the dish. Embryos at the desired stages were chosen and imaged using an Apochromat 20x/0.8 NA objective on a Zeiss LSM 880 confocal at 5 min intervals.

#### Development with auxin depletion

Embryos were collected from a self-cross of *NIPBL-mAID-mCherry*; *nanos>OsTIR(F74G)*/*His2Av-GFP* flies for 2 hours and were processed as described above and mounted and immersed in either DMSO or auxin supplemented 1x DPBS. Embryos were imaged every 15 min for 12 hours using an Apochromat 20x/0.8 NA objective on a Zeiss LSM 880 confocal.

### Transcription live imaging

#### Auxin depletion

Stable stocks of *NIPBL-mAID-mCherry*; *nanos>OsTIR(F74G)* were crossed with *NIPBL-mAID-mCherry /CyO*; *MCP-GFP, NLS-PCP-mCherry,His-BFP (TR)/TM6b. NIPBL-mAID-mCherry*; *nanos>OsTIR(F74G)/TR* females from the cross were selected and crossed to the requisite MS2 and PP7 tagged flies. Embryos were collected from the resulting cross for 90-120 min, dechorionated and incubated in 1x DPBS containing either 0.25 % DMSO (Control) or 50 μM 5-PH-IAA-AM (ΔNIPBL) for 90 min.

For the no OsTIR control (**Supplementary Fig. 4C**), *nanos>SV40NLS-PCP-mCherry, His2Av-BFP/CyO; nanos>MCP-GFP* females were crossed to wildtype *scyl-MS2, chrb-PP7* flies. Control embryos were untreated before imaging and auxin treatment was performed as described above.

#### WAPL mutant

*wapl*^*2*^/*FM7c* flies were crossed to *NIPBL-mAID-mCherry*; *TR/TM6b* flies and *wapl*^*2*^/+; *NIPBL-mAID-mCherry /+;TR/+* females were selected from the progeny and crossed to the appropriate MS2 and PP7 tagged males. Embryos were collected from the resulting cross for 3 hours and dechorionated using 4.12 % sodium hypochlorite solution and washed thoroughly before mounting.

#### Imaging settings

The embryos were then mounted on a low-fluorescence semipermeable membrane (Lumox film, Sarstedt cat. #94.6077.317) using double-side tape glue dissolved in n-heptane. The embryos were then immersed in Halocarbon oil 27 (Sigma-Aldrich #H8773) and sealed with a coverglass (#1.5). The imaging was performed using a Plan-Apochromat 40x/1.3 NA DIC M27 objective on a Zeiss LSM 880 confocal using a two-track setup with PCP-mCherry on one track and MCP-GFP on another track. The selected field of view was continuously imaged using 21 z-stacks separated by 0.5 μm with pixel averaging window of 2 frames. Each time frame of the movie took 54.3 s. The His-BFP signal was used to align embryos at mitosis and the 405 nm laser was switched off for the duration of the analyzed movie to prevent photobleaching and reduce autofluorescence in the RFP channel.

#### Image analysis

The pipeline for live imaging analysis can be found on github (https://github.com/pavancss/LongRangeGeneRegulation/). The pipeline is described concisely below. *Preprocessing and Nuclear Tracking*

Raw 3D Z-stacks (0.5 μm intervals) were converted into 2D time-series using maximum intensity projections. Nuclear segmentation was performed on the gaussian-kernel smoothed red fluorescence channel using the Cellpose^6^ convolutional neural network architecture. Nuclei were tracked across time-series based on substantial overlap across subsequent time points and nuclei tracked for less than five timepoints or intersect with the edge of the field of view.

##### Transcription Foci Segmentation

Transcription foci were identified within tracked nuclear domains using a local signal-to-background thresholding strategy. For each nucleus, the local background was estimated using Otsu’s thresholding of the intranuclear space. Active transcription sites were defined as contiguous pixels of a minimum size where the intensity ratio exceeded empirical thresholds determined by signal-to-noise optimization (6 for MS2; 2.5 for PP7).

##### Quantification of Transcriptional activity

Nuclear activity in both MS2 and PP7 channels was recorded as a function of time. Nuclei that showed a non-zero *scyl* transcriptional activity were marked as *scyl* positive nuclei. Since the *scylla* gene is located proximal to the enhancer, it was used as a marker for nuclei in which the enhancer was active. Fluorescence intensity traces for these nuclei were then analyzed to generate both an instantaneous and the cumulative *chrb* activity as a fraction of these nuclei. The total transcription in these nuclei was measured as the background subtracted fluorescence intensity of all the bursts at any given time point in all the *scyl*-positive nuclei. All the embryos imaged for each condition were aligned to the end of the movie (beginning of gastrulation) and the intensities/nuclear fraction values for all the embryos at every aligned time point were averaged to obtain the mean and standard error. This was done only for time points where a minimum of three embryos were imaged to minimize the variance.

##### Bursting analysis

Bursts were defined as continuous periods of non-zero nuclear activity. Bursts were labeled as distinct if the nucleus showed at least one frame with no activity in between. *chrb* bursts were only analyzed in nuclei that show *scyl*-positive nuclei as defined before. The time span of each burst defined as the ON durations and the time between bursts, defined as the OFF durations were extracted from the nuclei that show *chrb* bursts. These durations, with annotations for right truncation at the end of imaging period, were collected from all the embryos for a given condition and then fitted to a parametric log-normal accelerated failure time model to derive the true ON and OFF duration distributions. This was performed using the LogNormalAFTFitter function from the lifelines library (https://lifelines.readthedocs.io/en/latest/fitters/regression/LogNormalAFTFitter.html). The burst sizes were defined as the integrated intensity of the focus in a single burst, iff the burst ended before the beginning of gastrulation.

### DNA-FISH

#### Sample preparation

Embryos were dechorionated and fixed with 4% formaldehyde in 1x PBS and 50% heptane for 20 minutes at room temperature prior to the protocol. Embryos were tripled washed with 1x PBS, 0.1% Tween 20 and incubated with RNase A for 1 hour at 37 C. Following this, embryos were sequentially washed in increasing concentrations (20%, 50%, 80%, 100%) of FISH wash buffer (35% formamide, 4x SSC, 0.1% Tween-20). Probe hybridization occurred after incubation in 100% FISH wash buffer, with ~ 500 ng of each probe in 120 µL hybridization buffer (50% formamide, 10% dextran sulfate, 5x SSC, 100 ng Herring Sperm, 0.1% Tween-20) at 88 C. The temperature was slowly decreased down to 37 °C over 2 hours and was left overnight. The following day, the samples were washed in FISH wash buffer over 2 hours, triple washed with PBS, 0.1% Tween 20, and treated with DAPI.

#### Imaging

Embryos were mounted in Prolong Diamond at cured overnight at room temperature prior to imaging. Imaging was done with a Zeiss Confocal LSM880, with a Zeiss Plan-Apochromat 40X/1.3NA DIC M27 oil immersion objective lens (used with Immersol 518F oil). Images were acquired with 3 separate channels (405 nm, DAPI; 560 nm, ATTO565; 630 nm, ATTO633) with 0.1 µm * 0.1 µm * 0.25 µm voxel dimensions, with 2x averaging and 16 µs pixel dwell over 40 z-stacks.

#### Image analysis

Images were analyzed with custom code to extract spot pairs and measure 3D distances. A DAPI counterstain was applied to segment nuclei utilizing cellpose^6^ to determine eligible pairs of point source functions (PSFs). Candidate spot pairs were identified as the closest pair of spots within the same nuclei. For each candidate spot, a 3D-gaussian was fit and localized to subpixel precision; 3D-distances were calculated from the centroids of the 3D-gaussians. A chromatic aberration correction was applied on the basis that the vector between a spot pair of two different colors in the same nucleus has a zero mean in each dimension. Outliers due to false spot segmentation showing unreasonably high distances were removed.

### Polymer Simulations

Loop extrusion simulations were performed building on the “polychrom” package from the “open2c” GitHub project (https://github.com/open2c/polychrom), which was developed from earlier polymer models of loop extrusion written by the Mirny lab^7,8^, and powered by the GPU accelerated molecular simulation toolkit openMM (https://github.com/openmm/openmm). All simulations were run on either an NVIDIA Tesla V100 or GeForce RTX 3080 Ti card. Polychrom uses a Langevin approach to simulate the dynamics of a polymer under several user defined energy constraints. Complete details of the energetic constraints and other parameters are specified in the python simulation scripts included in our GitHub repository for this project: https://github.com/tee-udon/boettiger_livecell.

To simulate chromatin dynamics mediated by loop extrusion, we first simulated the location of cohesin left and right arms along the chromatin chain using 1D lattice model over time then we input the locations of these extruders into a molecular dynamics simulation to simulate the looping between pieces of the genome. We then saved snapshots of 3D trajectories to study the dynamics of chromatin structure over time.

Cohesin kinetics were simulated using a kinetic Monte Carlo (kMC) algorithm to capture stochastic binding, unbinding, and translocation events. The model starts with N_cohesin, unbound to the polymer. In each timestep, unbound cohesin complexes associated with the chromatin chain with a probability determined by the characteristic lifetime variable, *cohesin_unbound_lifetime*, while bound complexes dissociated with a probability derived from *cohesin_bound_lifetime*. Loading probabilities varied by genomic position as specified in the loading profile, *cohesin_loading_probability_list*. While binding, cohesin extruded loops by translocating bidirectionally at a rate of V_extrusion per timestep per arm.

When a translocating cohesin encountered another bound complex, it stalled with a probability, *cohesin_stall_probability*, and remained paused for an average duration of *cohesin_stall_lifetime* before resuming motion or dissociating. Additionally, CTCF binding sites were defined by their monomer coordinates and directionality (*ctcf_site_location_list, ctcf_direction_location_list*). Interactions between cohesin and CTCF sites with appropriate orientation stalled extrusion with probability *ctcf_site_stall_probability_list* for an average duration of *ctcf_site_stall_lifetime_list*. The 1D positions of all cohesin arms were recorded after every timestep for downstream MD integration.

Tether effects were simulated as sticky interactions between monomers, modeled with a Leonard-Jones type square well potential energy, as described previously. Monomers within the distance *attraction_radius* experience negative (favorable) potential energy, which transitions back to zero outside of the radius with a smooth, differentiable function, which helps ensure numerical stability. Monomer states were specified by the input *monomer_type_list*. Each monomer in type in the list then has a potential sticky interaction with the other types as defined by the input *attraction_matrix*. We used a simple two state system, where all monomers are type 0 except tethers are type 1, and only type 1 experience homotypic weak interactions 1.5 kT.

The chromatin fiber was initialized as a polymer in a spherical fractal conformation. The 3D conformational changes were dynamically coupled to the 1D lattice model through an iterative update scheme. For each timestep of the extrusion simulation, the coordinates of the left and right cohesin arms were mapped to the polymer chain. These positions were used to impose transient pairwise harmonic bonds between the corresponding monomers, effectively simulating the formation of chromatin loops. Following the update of these looping constraints, the polymer system was evolved for a duration of *num_MD_steps_per_LE* to allow for local relaxation and structural reorganization. This cycle was repeated for the duration of the simulation, with 3D structural snapshots recorded at defined intervals to characterize the temporal evolution of the chromatin architecture.

Transcriptional activation was modeled as a stochastic process, dependent on the amount of time the enhancer spent in proximity to the promoter. When the 3D distance between the enhancer monomer and the promoter monomer are less than *enhancer_promoter_contact_threshold*, they are considered in proximity. The probability of a promoter firing is determined by the promoter activity. Promoter activity increased by one unit per time step while the enhancer and promoter are in proximity. The model assumes this process is dominated by a single rate limiting step, so the increase occurs after an exponentially distributed waiting time, *promoter_activation_time*. The promoter activity then drops back by one unit after exponentially distributed waiting time, *promoter_deactivation_time*. This ensures that each contact leads to an increase in promoter state, but without continued contact the promoter state returns to its resting 0. Finally, to record a transcript, the promoter state must reach state *promoter_state_fire_threshold*. A value of 1 means even the briefest kiss-and-run contact will result in a transcript, whereas larger values require more frames of contact in order to elicit a response.

The parameters used for the simulations condition presented in the text are detailed below. The same parameter value was used in all simulations, except as indicated below - where individual parameters were changed to simulate the effects of specific perturbations.

*num_monomers* = 1500

*num_cohesin* = 5 (*num_cohesin* = 0 for NIPBL degron simulations)

*cohesin_unbound_lifetime* = 1

*cohesin_bound_lifetime* = 200 (*cohesin_bound_lifetime* = 400 for WAPL het. simulations)

*cohesin_loading_probability_list* = uniform

*cohesin_speed* = 1

*cohesin_stall_probability* = 1

*cohesin_stall_lifetime* = inf

*ctcf_site_location_list, ctcf_direction_location_list*. = 500, bidirectional.

*ctcf_site_stall_probability_list* = 1 (0 for no CTCF simulations)

*ctcf_site_stall_lifetime_list* = 1000

*Monomer_type_list* = [[0,492],[1,8],[0,485],[1,15],[0,500]]

*Attraction_coefficient_matrix* = [[0,0],[0,1.5]] ([[0,0],[0,0]] for no tether simulations).

*attraction_radius* = 1.5

*num_MD_steps_per_LE* = 500

*enhancer_promoter_contact_threshold* = 9

*promoter_activation_time* = 500

*promoter_deactivation_time* = 50

*promoter_state_fire_threshold* = 8

Monomer 500 was designated the enhancer and monomer 1000 the chrb promoter. The extra additional 500 monomers upstream and downstream (1500 total) were added to avoid edge effects.

Large language models (ChatGPT, Claude and Gemini) were used for assistance with manuscript preparation. All scientific content, data analysis, and conclusions were verified by the authors.

### L.L.M Usage

Large language models (ChatGPT, Claude and Gemini) were used for assistance with manuscript preparation and code generation. All scientific content, data analysis, and conclusions were verified by the authors.

